# Weak Coupling Between Spontaneous Local Cortical Activity State Switches Under Anesthesia Leads to Strongly Correlated Global Cortical States

**DOI:** 10.1101/2021.11.07.467591

**Authors:** Ethan B. Blackwood, Brenna P. Shortal, Alex Proekt

## Abstract

Under anesthesia, neural dynamics deviate dramatically from those seen during wakefulness. During recovery from this perturbation, thalamocortical activity abruptly switches among a small set of metastable intermediate states. These metastable states and structured transitions among them form a scaffold that guides the brain back to the waking state. Here, we investigate the mechanisms that constrain cortical activity to discrete states and give rise to abrupt transitions among them. If state transitions were imposed onto the thalamocortical system by changes in the subcortical modulation, different cortical sites should exhibit near-synchronous state transitions. To test this hypothesis, we quantified state synchrony at different cortical sites in anesthetized rats. States were defined by compressing spectra of layer-specific local field potentials (LFPs) in visual and motor cortices. Transition synchrony, mutual information, and canonical correlations all demonstrate that most state transitions in the cortex are local and that coupling between sites is weak. Fluctuations in the LFP in the thalamic input layer 4 were particularly dissimilar from those in supra- and infra-granular layers. Thus, our results suggest that the discrete global cortical states are not imposed by the ascending modulatory pathways but emerge from the multitude of weak pairwise interactions within the cortex.

## Introduction

Brain activity arises as a result of interactions amongst billions of neurons and synapses. Each component in this vast network exhibits complex nonlinear dynamics (Hodgkin and Huxley, 1952; Pan and Zucker, 2009). Generically, such complex nonlinear dynamical systems can dramatically change their collective behavior after small changes in parameters or perturbations to their ongoing activity (Canavier et al., 1993; Destexhe et al., 1994; Ermentrout, 1998; Izhikevich, 2007; Strogatz, 2015). Furthermore, because nonlinear systems generally have multiple steady state behaviors, there is no guarantee that after a dramatic perturbation, the system will recover to its previous state once the perturbation subsides.

These considerations suggest that brain activity ought to be quite fragile and unable to withstand dramatic perturbations. Contrary to this intuition, there is ample evidence that the brain is remarkably robust to perturbations. Seizures, for instance, are a paradigmatic example of aberrant brain activity, being characterized by extreme synchronization in neuronal firing and subthreshold voltage fluctuations (Timofeev et al., 2004). While seizures can be followed by a transient postictal period characterized by abnormal brain activity and function (Fisher and Engel, 2010), normal brain function is eventually restored. Another classic example of the brain’s ability to recover from an extreme perturbation is general anesthesia (Brown et al., 2010). Every year, millions of patients undergo general anesthesia. While some patients experience aberrant brain activity, which manifests as delirium upon emergence (Saczynski et al., 2012), most eventually recover normal brain activity and cognitive function. During general anesthesia, the brain may exhibit dramatically abnormal activity patterns, such as burst suppression, which is caused by the hyperpolarization and silencing of more than 90% of cortical neurons (Amzica, 2009; Civillico and Contreras, 2012; Contreras and Steriade, 1997). Occasionally, complete isoelectric electroencephalogram (EEG) is observed in surgeries requiring circulatory arrest (Stecker et al., 2001). Nevertheless, once anesthetic delivery is stopped, the brain regains normal function. Given this and the fact that anesthetic delivery can be precisely controlled, general anesthesia is a good model system to address the general question of how the brain is able to restore normal activity patterns after a dramatic perturbation.

Several converging lines of evidence strongly argue that recovery from anesthesia cannot be explained by anesthetic washout alone. The first is that recovery of consciousness after anesthesia occurs at a lower anesthetic concentration than induction of anesthesia across taxa, from *Drosophila* (Joiner et al., 2013) to mice (Friedman et al., 2010) and humans (Warnaby et al., 2017). Furthermore, this neural inertia can be modulated by factors altogether unrelated to the concentration of anesthetic, such as single gene mutations (Friedman et al., 2010) and manipulations of specific neuronal populations (Kelz et al., 2008; Reitz et al., 2021; Zhou et al., 2018). Together, these results strongly argue that recovery from anesthesia is not simply the byproduct of anesthetic washout. They do not, however, directly shed light on the mechanisms that allow the brain to recover after general anesthesia.

In order to recover from anesthesia, the brain must follow a path through the state space that begins in the deeply anesthetized state and eventually leads back to the pre-anesthetic conditions. The neurophysiological processes that allow the brain to navigate this path efficiently have been addressed by Hudson et al. (2014). Specifically, they show that *en route* to recovery of consciousness, brain activity is constrained to a low-dimensional space. In this space, most activity is confined to a small number of discrete activity patterns, and the transitions between these patterns are highly structured. In sum, these mechanisms greatly constrain the number of possible paths through the activity space that can lead to wakefulness and allow the brain to recover consciousness on a physiological time scale. Abrupt transitions between discrete activity states have been observed in rodents (Hudson et al., 2014), non-human primates (Ballesteros et al., 2020; Ishizawa et al., 2016; Patel et al., 2020) and human patients (Chander et al., 2014) after exposure to a variety of anesthetics with distinct mechanisms of action. Abrupt transitions between different activity patterns at a fixed anesthetic concentration are observed not only at the level of the local field potentials (e.g., Hudson et al., 2014), but also in the activity of individual cortical neurons (Lee et al., 2020). These discrete activity patterns and structured transitions between them serve as a scaffold that guides the brain back towards normal patterns of activity after it has been profoundly disrupted by anesthetics.

Given that state transitions are critical for reinstating consciousness, it is of fundamental importance to determine the neuronal mechanisms that give rise to transitions between discrete activity states during recovery from a dramatic perturbation. Previous work on anesthesia (Chander et al., 2014; Hudson et al., 2014; Ishizawa et al., 2016) and sleep (Gervasoni et al., 2004) defined different activity patterns on the basis of oscillatory activity observed in the local field potentials (LFPs) of firing of individual neurons (Lee et al., 2020). Much of this oscillatory activity is coordinated via thalamo-cortical loops (Contreras and Steriade, 1997; Liu et al., 2015; Schiff, 2008; Steriade et al., 1993b). An extensive body of work shows that the thalamocortical circuitry is modulated by the arousal pathways ascending from the brainstem and basal forebrain to produce oscillations at different characteristic frequencies (Destexhe et al., 1994; Jones, 2003; Steriade et al., 1993a). Indeed, during constant anesthetic concentration, fluctuations in the firing rates of individual neurons within these arousal nuclei co-vary with fluctuations in the spectra of cortical LFPs (Gao et al., 2019). Direct manipulations of neuronal activity within the reticular activating system can elicit profound changes in the oscillations observed in the cortical LFP (Gao et al., 2019; Moruzzi and Magoun, 1949; Steriade et al., 1993a; Vazey and Aston-Jones, 2014). Thus, one distinct possibility is that the discrete oscillatory patterns of activity observed under fixed anesthetic concentration are imposed onto the thalamocortical networks by fluctuating modulatory tone. If this is the case, because modulatory systems project broadly across the thalamus and cortex (Jones, 2003), we expect to find that abrupt transitions between distinct oscillations occur in close temporal proximity across the different cortical layers and regions. Alternatively, it is possible that the oscillatory activity in different cortical regions is largely coordinated through short-range thalamo-cortical and cortico-cortical interactions. In this case, we expect to find that transitions between different oscillatory patterns are largely local.

Here, we provide direct experimental evidence for this latter possibility by simultaneously recording abrupt transitions between different states across cortical layers and across distant cortical areas at a constant anesthetic concentration. Using a complementary combination of analytic techniques, we show that state transitions across different cortical sites are only weakly coupled. Furthermore, we demonstrate that state transitions in layer 4 (L4)—the layer that directly receives input from the thalamus—are particularly decoupled from state transitions observed in other layers. This suggests that cortico-cortical interactions rather than fluctuations in the broad modulatory tone play a crucial role in controlling state transitions under anesthesia. Remarkably, we also show that the multitude of weak pairwise interactions between local state transitions is sufficient to constrain the overall brain activity to just a few states embedded in a low-dimensional space. Thus, our results suggest that the highly coordinated, low-dimensional macroscopic brain dynamics that allow the brain to recover from a dramatic perturbation emerge as a consequence of a multitude of weak pairwise interactions between different cortical sites.

## Materials and Methods

### Animals

All experiments were performed using ten male Sprague-Dawley rats, each two to three months of age (250–350 g) (Charles River Laboratories, Wilmington, MA). Two animals were excluded from further analyses because of excessive burst suppression or noise, respectively. One additional animal was excluded after current source density analysis revealed that the V1 probe was inserted too deeply to clearly identify cortical L4 and the supragranular layers. Rats were housed under a conventional 12:12 h, light:dark cycle and given food and water *ad libitum*. All experiments were performed in accordance with the Institutional Animal Care and Use Committee at the University of Pennsylvania and conducted in accordance with the National Institute of Health Guidelines.

### Surgery

All surgeries were performed under aseptic conditions. Each animal was weighed immediately prior to surgery. Animals were induced with 2.5% isoflurane in oxygen and secured in a stereotaxic frame (Kopf Instruments, Los Angeles, CA) in the prone position. Core body temperature was maintained at 37 (± 0.5) °C using a temperature controller (TC-1000 Temperature Controller, CWE, Incorporated, Ardmore, PA). Prior to surgery, isoflurane concentration was reduced to 1.5% (flow rate 1 L/min), and dexamethasone (0.25 mg/kg) was delivered subcutaneously. Bupivacaine (5 mg/mL) was injected under the scalp to provide local anesthesia. Throughout the surgery, the lack of response to a toe pinch was used to assess proper anesthetic depth.

The scalp was retracted and two 2 × 2 mm craniotomies were performed using a dental drill: one centered over -5.52 mm AP, 4 mm ML of bregma and another centered over -1.26 mm AP and 1.55 mm ML of bregma for V1 and M1 sites respectively. Dura was removed and Gelfoam (Pfizer, New York, NY) was placed on the exposed cortical tissue to prevent the tissue from desiccating. Prior to insertion, both linear probes (Cambridge NeuroTech, Cambridge, UK; H3 acute 64-channel linear probe) were dipped in DiI to allow for subsequent track tracing and lowered to 1.2 mm into the brain. Prior to electrode insertion, Dura Gel (Cambridge NeuroTech) was applied to each craniotomy and isoflurane concentration was lowered again to 1% (flow rate 1 L/min) for recordings. Immediately following electrophysiological recordings, animals were perfused trans-cardically with 4% paraformaldehyde under 4% isoflurane. Brain was harvested and processed for electrode track tracing.

### Histological confirmation of recording sites

Brains were sectioned at 80µm on a vibratome (Leica Microsystems, Wetzlar, Germany). Sections were mounted with medium containing a DAPI counterstain (Vector Laboratories, Burlingame, CA). Electrode tracks were manually identified and localized using epifluorescence microscopy (Olympus, Tokyo, Japan; BX41) at 4x magnification.

### Electrophysiology and Preprocessing

All recordings were performed at 1% isoflurane, after allowing the anesthetic concentration to equilibrate for at least 30 minutes. Signals were amplified and digitized on an RHD2132 headstage (Intan, Los Angeles, CA) and streamed to a PC using an Omniplex acquisition system (Plexon, Dallas, TX) at a rate of 40,000 samples per second per channel. All recordings were performed using a ground skull screw as reference. Local field potentials (LFP) were extracted from raw signals online using the bandpass filter with a passband of 0.1-300 Hz. Offline, LFP were decimated to 1 kHz and filtered using a custom acausal FIR 0.1–200 Hz bandpass filter. Noisy channels were removed by visual inspection of the signals. Before subsequent analyses, data were re-referenced to the mean computed over all clean channels on the laminar probe. All data analysis was completed using custom built MATLAB (MathWorks, Natick, MA) code unless otherwise stated. In total, 29.88 hours of recordings were used to generate all data in this manuscript.

### Current Source Density and Channel Selection

In order to facilitate cortical layer localization, a series of 10 ms light flash stimuli was presented from a green LED positioned about one inch from the eye contralateral to the craniotomy over V1. Interstimulus intervals were drawn from a uniform distribution between 3 and 5 seconds to prevent stimulus entrainment. Current source density (CSD) analysis was then applied to the post-stimulus LFP to identify layers in V1. The CSD C_t_ at time *t* was calculated by computing a smoothed second spatial derivative (a representative example is shown **Figure 5**):

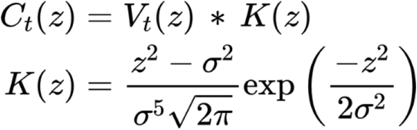

Here, *z* is the channel depth, σ = 280 μm is the distance along the electrode from *z* at which the kernel changes sign, *V*_*t*_ is the mean voltage over all light flash trials at time *t* relative to flash onset, and ∗ indicates convolution. The electrode closest to the center of L4 was identified manually from the CSD as the earliest current sink. Once L4 was identified, supra- and infragranular channels were selected for analysis at 140 μm intervals above and below L4.

### Time-Frequency Analysis

Spectrograms of selected channels were calculated from LFP signals using the multitaper method with 17 Slepian tapers and time-bandwidth product (NW) = 9. A 6-second sliding window with a step size of 100 ms was used. Windows containing signal artifacts were identified and removed using a combination of automatic burst suppression detection based on the root-mean-square of LFP in a moving exponential window and manual inspection of multitaper spectrograms. Each window was zero-padded to 65.536 s to increase the frequency resolution and input a power-of-2 number of samples to the Fourier transform. In order to sample frequencies of greater interest more densely, 279 frequencies were selected from 0.14 to 300 Hz, spaced on a log scale from 0.14 to 10 Hz and on a linear scale above 10 Hz. The multitaper spectrograms were then smoothed over frequencies with a median filter spanning 10 frequency steps (up to 17.5 Hz) and over time with an exponential (Poisson) window spanning 2 minutes. In order to remove baseline differences in power across frequencies (such as power-law scaling) and emphasize temporal fluctuations, each spectrogram was rank-order normalized along the time axis. At each frequency bin, the time window with the highest power was given the value of one. Each other window was given the value of (*r*−1)*/*(*N*−1), where *r* is that window’s sorted index among the *N* windows. Thus, the smallest power value at each frequency was represented as zero, and the largest as one.

### Dimensionality Reduction

Dimensionality reduction was performed on each channel’s spectrogram individually, in order to obtain high reconstruction accuracy and ensure that any characteristic differences in activity patterns between sampled regions and cortical depths were preserved. Non-negative matrix factorization (NMF) (Lee and Seung, 1999; Mankad and Michailidis, 2013) was used to compress the rank-ordered spectrograms. The NMF output represents the signal at each time as a short vector of *K* non-negative coefficients (scores) that weight a sum of corresponding frequency components (loadings) to reproduce the original spectrum. Given a spectrogram *A* of size 279 x *N*, NMF produces a loading matrix *U* of size 279 x *K* and a score matrix *V* of size *N* x *K*. The product *UV*^T^ reconstructs *A* with some error *E*, quantified relative to the norm of *A* as:

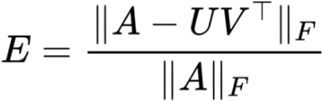

Where ‖·‖_*F*_ is the Frobenius norm. To select an appropriate number of components (*K*) for each channel, a cross-validation approach was employed (Owen and Perry, 2009). First, spectrograms were downsampled across time by a factor of 20, for computational efficiency. Then, a random subset of 20% of the rows and columns were selected to be withheld. Starting with *K* = 1 and increasing to 15, NMF was applied to the down-sampled matrix after the random subset of rows and columns had been removed. This iteration provides both a loading and score matrix. Next, NMF was run again on the data with only the pre-selected rows withheld. In this iteration, the loading matrix from the first round was fixed and only a new score matrix was calculated. In the third and final run of NMF, NMF was run on the data with only the pre-selected columns removed, fixing the score matrix from the first round and calculating only a new loading matrix. Finally, the loading and score matrices produced in the second and third run of NMF, respectively, were multiplied to generate an estimate of the original dataset and calculate error as a function of *K*. This procedure was repeated for five replicates for each value of *K*, and the optimal *K* was chosen such that increasing *K* by one would reduce mean reconstruction error by less than 1%. In our dataset, the optimal value for *K* ranged from five to nine for different channels. After the cross-validation procedure, each channel’s full, normalized spectrogram was subjected to NMF using the channel’s optimal *K*, resulting in a mean reconstruction error of 14.8% across all channels (∼85% of the variance captured by NMF for each spectrogram). Note that NMF does not constrain the relative scales of the loading vectors: for any invertible diagonal *K* x *K* matrix *D, UV*^T^ = *UD*(*VD*^-1^)^T^. To remove these degrees of freedom, *U* and *V* were rescaled by a matrix *D* such that the rescaled loadings had unit L_2_ norm.

### Transition and Discrete State Identification

The rescaled score matrix *VD*^-1^ is the basis for defining each channel’s state over time. For each channel, at each time point, the component with the highest score was taken as the state of the brain near that channel’s recording site, and samples where the state changed were marked as local transition times. In order to prevent an arbitrarily high number of transitions during periods when two or more components had similar scores, transitions that were likely to reflect transient fluctuations were ignored and the state assignments between them were updated accordingly. Specifically, suppose one time segment between two transitions was assigned state “A” and either the previous or next segment was assigned state “B.” If the first segment was less than 100 seconds long and, within the first segment, the mean score for NMF component A was less than 1.1 times the mean score for component B (*i*.*e*., if the state assignment was sufficiently ambiguous), the transition between the two segments was ignored and the combined segment was assigned state B. If a segment could be merged with either the previous or next segment, the tie was broken by ignoring the transition with a smaller magnitude of change in the full NMF score vector from the 3 seconds before the transition to the 3 seconds following it. A matrix of state transition frequencies was computed by tabulating how often each discrete state followed each other state over the duration of the recording using the table of discrete state transitions for each channel.

### Markov-based Shuffled Null Model

When testing whether pairs of channels are synchronized in the sense that they preferentially occupy certain combinations of discrete states, apparent synchrony could arise due to the channels’ individual NMF score distributions, independent of the relative timing of transitions. To control for this possibility, a discrete-time Markov chain (the “null model”) was fit to the transition frequencies of each channel independently. The channel’s null model was then used to simulate 1000 new discrete state sequences of the same length as the original data. For each pair of channels, these “null” state sequences were then used to fit distributions of transition synchrony and normalized mutual information (see corresponding sections below). This distribution reflects the probability of observing a given state synchrony and mutual information under the assumption of complete independence between different recording sites. To obtain a null distribution of canonical correlation-based synchrony (see below), full score matrices were generated from each channel’s null state sequences as follows: for each of the *K* states *k*, at each sample with null discrete state assignment *k*, the corresponding row of the null score matrix was randomly drawn from the set of rows of the original data score matrix where the original discrete state was equal to *k*. These random sequences for all pairs of channels were then subjected to canonical correlation analysis.

After fitting normal distributions for each of the three channel pair interaction measures (transition synchrony, normalized mutual information, and canonical correlations) to the shuffled surrogates, the values obtained for the real data were tested against these distributions to estimate whether they would be expected by chance, given the statistics of the data (see “Statistical Tests” below).

### Transition Synchrony

To quantify how frequently channels transitioned together we employed the SPIKE-synchronization score (“synchrony score”), a method for quantifying synchrony between two simultaneously recorded sequences of events (Kreuz et al., 2015). At its core, this method is a coincidence detector in which the coincidence window is derived from the dataset. The adaptive definition of the coincidence window means that this method for quantifying synchrony is equally well-suited for state transitions as it is to spike trains. Each transition *r* is assigned a local window length τ(*r*), which is defined as half the smaller of the inter-transition intervals directly before and after *r*. For a pair of channels *i* and *j*, if transition *r*_*j*_ in *j* was the closest transition to transition *r*_*i*_ in *i* and vice versa, and the time between *r*_*i*_ and *r*_*j*_ is less than min(τ(*r*_*i*_), τ(*r*_*j*_)), both transitions have a synchrony score of 1. All other transitions have a score of 0. This measure is extended to the multi-channel case by assigning each transition a synchrony score equal to its mean pairwise synchrony score with the nearest transitions in all other channels. Both pairwise and all-channel synchrony scores were computed for all discrete state transitions in each recording, and then averaged over all transitions to obtain pairwise and global mean synchrony measures.

### Normalized Mutual Information

Mutual information of discrete states was used to quantify the synchrony of states themselves rather than just the timing of their transitions. Specifically, this measure was implemented to quantify how well one could predict the state in one channel, given the state of another channel at the same time point. Since NMF was performed separately on each channel, states labeled with the same index in different channels are not necessarily the same with respect to the frequency characteristics of the signal. Regardless, mutual information is able to reveal temporal relationships between channel pairs because it does not assume any particular relationship between the state assignments of the different channels and is, therefore, agnostic to the assignments themselves.

Mutual information *I*(*X; Y*) between two channels *X* and *Y* with *N* observations and sets of classes *K*_*X*_ and *K*_*Y*_ was computed pointwise as follows:

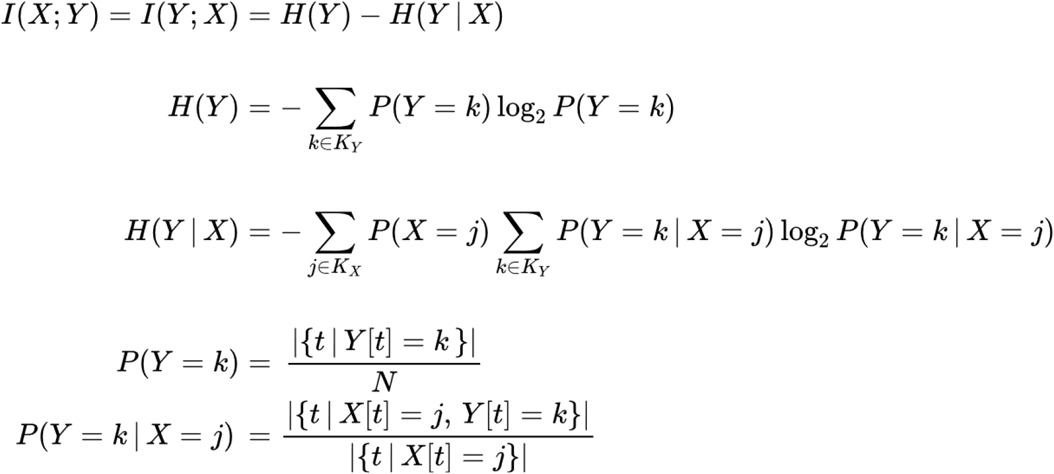

Mutual information is not a pure measure of the predictability of one variable given the other; it also increases with the entropy of each variable. For example, if channels *X* and *Y* each occupy a wider distribution of states and, as a result, have higher entropy than both channels *W* and *Z*, then *I*(*X*; *Y*) > *I*(*W*; *Z*). This is true even if the state of *X* is perfectly predictable given *Y, Y* given *X, W* given *Z*, and *Z* given *W*. In order to control for this, mutual information was normalized by the sum of the entropies of the two channels, giving the normalized mutual information, or symmetric uncertainty (Witten et al., 2011):

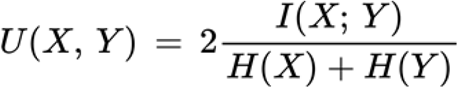

Using another definition for mutual information in terms of the individual and joint entropies of *X* and *Y*, we can write:

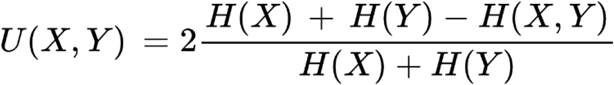

Thus, normalized mutual information can be understood as twice the fraction of the sum of individual entropies, *H*(*X*) + *H*(*Y*), that exceeds (is redundant to) the joint entropy *H*(*X, Y*) due to mutual information between *X* and *Y*. For example, if *X* and *Y* are identical, *U*(*X, Y*) = 1 and 50% of *H*(*X*) + *H*(*Y*) is redundant, as only one of the variables carries unique information.

### Canonical Correlation

Both the transition synchrony and normalized mutual information measures assume that LFP signals at each channel form discrete states and that the sequence of NMF components with the largest magnitude at each time point is informative about this state. However, there may be cases where multiple components must be considered. For instance, consider a situation in which NMF component A in channel *i* is characterized by strong activity in two frequency bands, and components B and C in channel *j* are characterized by strong activity in one of those frequency bands each. If only the “top” component determines the discrete state, there could be artificially low synchrony and mutual information between channels *i* and *j*. This is because, during a bout of state A in channel *i*, there could be frequent switching between states B and C in channel *j*, even though the overall signal characteristics in channel *j* remain largely static. To address this kind of ambiguity and compute a state synchrony measure that softens the artificially sharp boundaries between “discrete states,” canonical correlation analysis (CCA) was applied to the NMF score matrices of pairs of channels. Intuitively, CCA allows each score matrix to be linearly transformed to optimally match components between channels. CCA maximizes the correlations between the matched, transformed components. These correlations are used to derive a measure of state similarity.

The procedure for computing CCA-based synchrony is as follows: let *V* ∈ ℝ^NxL^ and *W* ∈ ℝ^NxM^ be the NMF score matrices two channels, and let K = min(L, M). At each step *i* from 1 to K, CCA finds coefficient vectors *a*_*i*_ and *b*_*i*_ to maximize the correlation *ρ*_*i*_ = corr(*Va*_*i*_, *Wb*_*i*_), with the constraints that *a*_*i*_ is uncorrelated with all previous vectors *a*_1_, …, *a*_*i*-1_, and likewise for *b*_*i*_. The MATLAB function *canoncorr* was used to perform this algorithm and the canonical correlation coefficients *ρ*_1_, …, *ρ*_*Κ*_ were averaged to obtain a state similarity measure.

### Statistical Tests

This section describes the procedure used to establish the statistical significance of interactions between recordings sites as measured by the synchrony score, normalized mutual information, and canonical correlation analysis. For each channel pair under consideration and each of these three interaction measures, the measure was computed both on the experimental dataset and on a set of 1000 null-model datasets generated from discrete Markov models of each channel’s transition statistics, as described above. The values of each measure were approximately normally distributed across null-model datasets. To test statistical significance, the deviation of each measure obtained in the experimental dataset from those generated from null-model datasets was expressed as a z-score. The one-tailed *p*-value was then directly computed from the z-score. The significance threshold was set at α=0.05. Bonferroni correction was applied to account for multiple comparisons over all channel pairs in each animal. The percentage of pairs for which each interaction measure was different from chance after Bonferroni correction is reported in the manuscript, and non-significant pairs are grayed out in Figures 6-8.

To compare interaction measures between different sets of channel pairs, special consideration must be paid to the statistical dependence between observations. In a recording with *n* channels, for any channel *k*, one would not in general expect the values of a distance-like measure on the pairs (*k*, 1), …, (*k, k-*1), (*k, k+*1), …, (*k, n*) to be independent. For example, if channel *k* were an outlier, all *n*-1 pairs would take extreme values due to what is statistically only one extreme observation. If pairwise statistics were compared naively, e.g., using a two-sample *t-*test, these dependencies would result in an overestimation of effective sample size and thus significance. Instead, a Monte Carlo permutation procedure was used to establish null distributions for comparisons of pairwise measures between groups of channel pairs. This procedure randomly shuffled group assignments while preserving the dependency structure inherent in the matrix of pairwise measures by only shuffling rows and columns. For each such comparison, 10^7^ permutations of only the channels of each recording that were included in that comparison were conducted, and the difference of group means was computed after each permutation. The frequency with which these null differences exceeded the difference of means of the unpermuted groups was taken as the *p-*value of the comparison.

Finally, when comparing the interaction measures for between-region channel pairs in M1/V1 recordings to those in bilateral V1 recordings, the method of permuting channel labels cannot be used because there are no data for pairs of channels that mix different recordings. Instead, the distribution for the difference of means of the measure over pairs between the two sets of recordings was estimated by bootstrapping over channels. Specifically, each group in such a comparison consists of a set of rectangular matrices, containing values of the measure for each pair of one channel along the rows and one channel along the columns. By resampling both rows and columns with replacement in each such matrix, the dependencies along rows and columns were preserved, but the variance in the mean could be estimated thanks to the principles of bootstrapping. A total of 10^6^ bootstrapped estimates of the group mean difference were computed in this manner for each interaction measure and used to obtain a *p-*value for the one-tailed hypothesis that the measure is greater on average between hemispheres of V1 than between M1 and V1.

## Results

### State transitions under constant anesthetic can be local

We sought to determine whether state transitions under a fixed concentration of isoflurane (1% atm.) occur simultaneously across different cortical regions and across layers within the same cortical region. This concentration was chosen based on previous work (Hudson et al., 2014) showing that burst suppression is not likely to occur at this concentration, but that state transitions in the spectral characteristics of the LFP are frequently observed. Here we focused on the local field potentials (LFPs) recorded using two laminar probes that sampled signals across all cortical layers. In half of the experiments, both electrodes were inserted into the right hemisphere: one in the primary visual area (V1) and the other in the motor cortex (M1) (n = 3) (**Figure 1A**). In the other half of experiments, bilateral V1 recordings were performed (n = 4). Postmortem localization of electrodes (Methods) in a representative experiment is shown in **Figure 1B**. Consistent with previous findings (Hudson et al., 2014), at 1% isoflurane, the power spectrum of the LFP fluctuated between several discrete states (**Figure 1C**).

**Figure 1:**
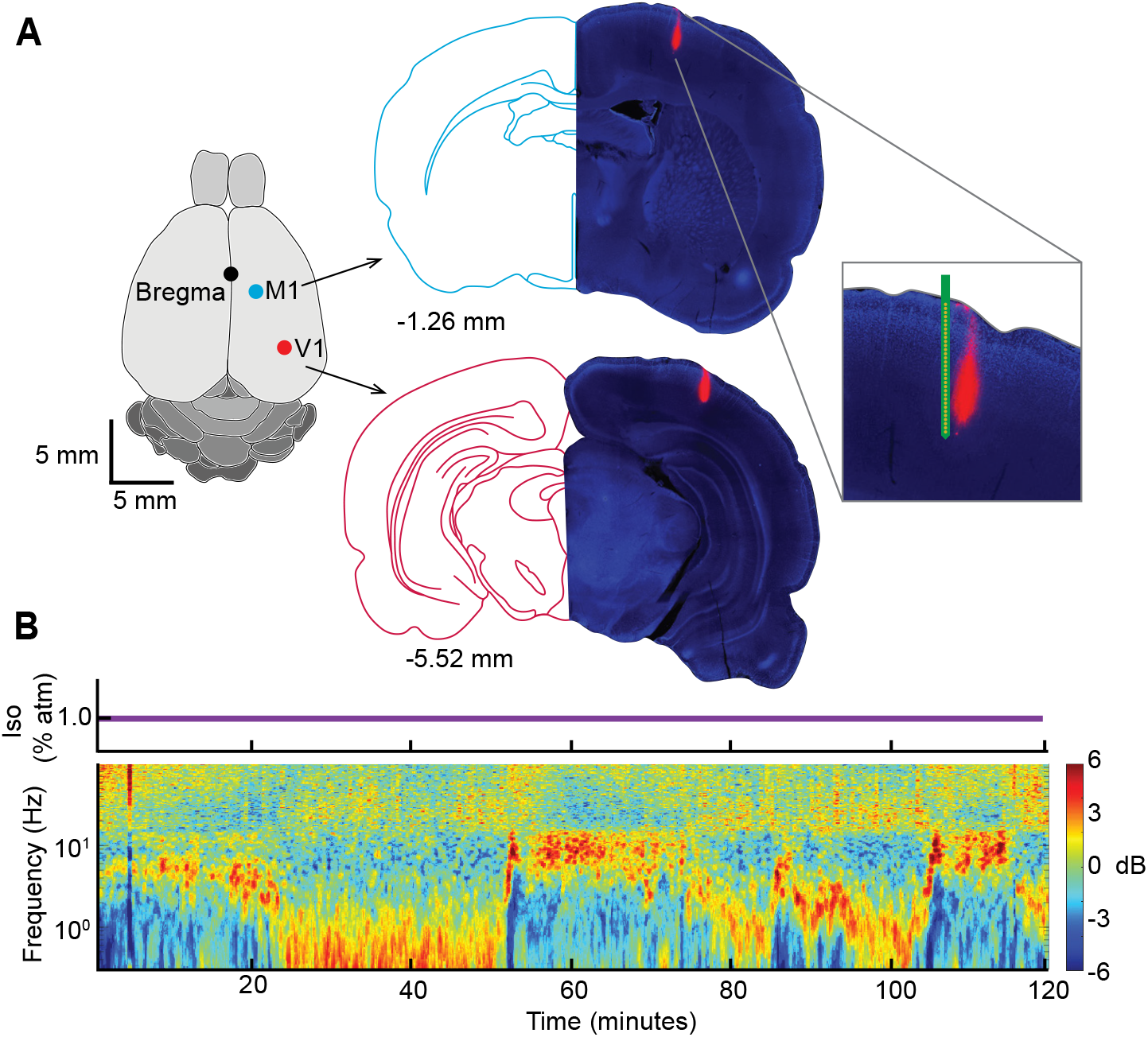
Experimental setup. **A**. Verification of Electrode placement into V1 and M1. DAPI-stained histological section showing tracks of the DiI-dipped electrode (right) juxtaposed with the corresponding section from the rat brain atlas (left). The zoomed cutout includes an image to show electrode channel layout. **B**. Time-resolved spectrogram recorded from V1 under 1% isoflurane general anesthesia (concentration shown above spectrogram). Spectrogram is plotted as deviations from temporal mean.

State transitions can be readily identified in the raw LFP (**Figure 2**). The top and bottom LFP traces show one minute of recordings from a single M1 and V1 electrode, respectively. The accompanying spectra were calculated using a multitaper spectral estimate. These spectra were averaged across two second windows of LFP with a one second step size, sampled either from eight to two seconds prior to transition (black, pre-transition) or from two to eight seconds after the transition (red, post-transition). Spectral estimates are shown as mean ± 95% confidence interval computed from 1000 bootstraps. In some instances, state transitions occur approximately simultaneously in the motor and visual cortices (**Figure 2A**). However, this was not always the case. For instance, **Figure 2B** shows an example of a state transition that occurs first in the visual cortex and, only after a delay of approximately 10 seconds, is seen in the motor cortex. Thus, abrupt changes in the LFP characteristics need not occur simultaneously in different brain regions. **Figure 2C** shows a more extreme example of this phenomenon. A state transition is clearly seen in the motor cortex, but in the visual cortex, the LFP characteristics remain unchanged. These observations suggest that, while some state transitions may indeed be global, there is a previously unappreciated degree of independence between state fluctuations observed in the cortex during fixed anesthetic administration.

**Figure 2:**
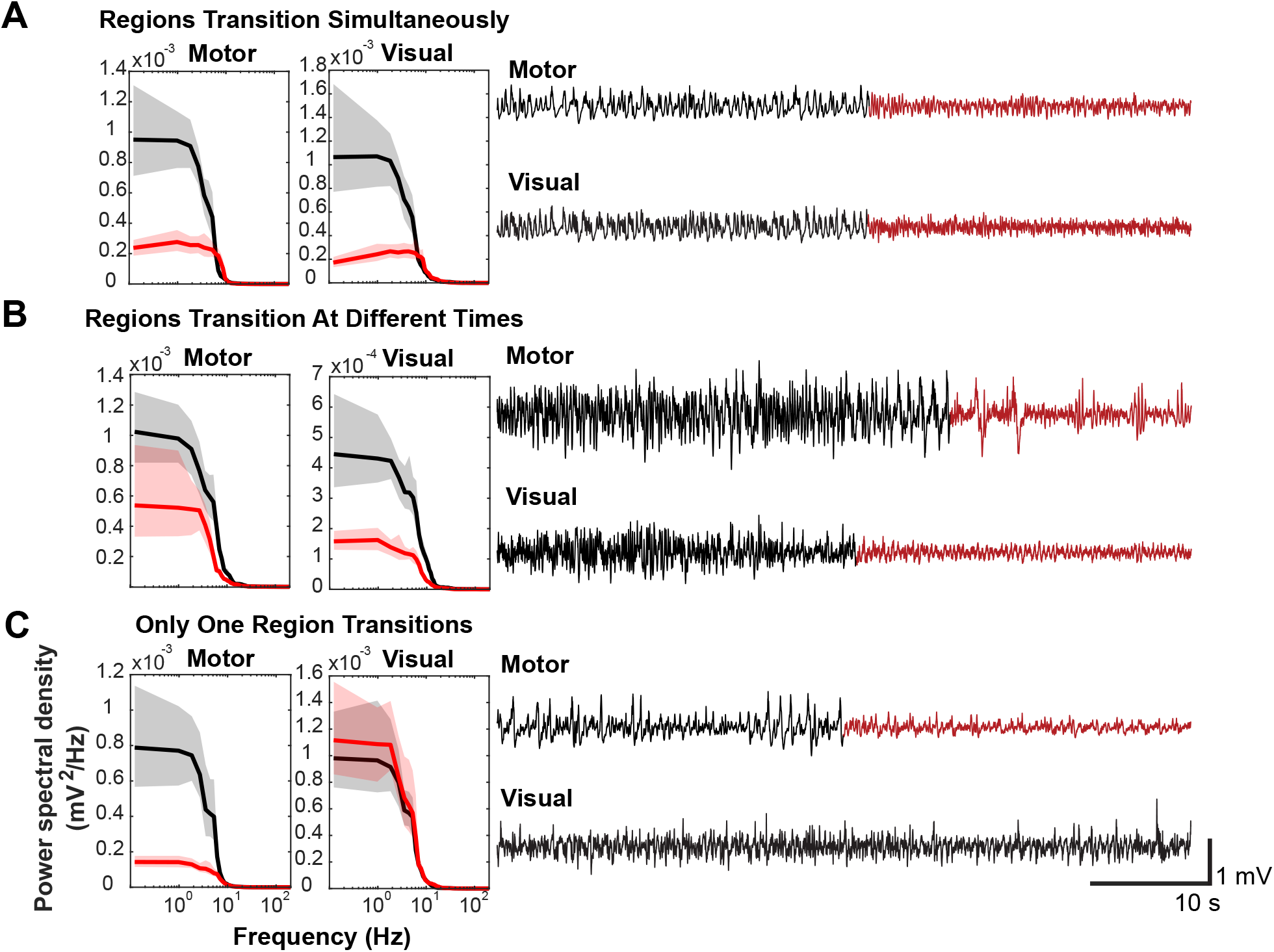
Examples of state transitions. **A-C** Right: LFP traces (1 minute) recorded simultaneously from right M1 and V1. Visually apparent abrupt transitions in the character of the LFP are indicated by shifts of color from black to red. Left: spectra computed from the red and black time periods respectively to indicate that the abrupt switches in the features of the signals are associated with changes in the spectra. **A**. An example where both M1 and V1 LFPs appear to change state simultaneously. **B**. An example where both M1 and V1 signals change states but with an appreciable time delay (∼10 s). **C**. An example where a state transition is observed in M1 but not in V1. In this case for the purposes of computing the spectrum (left, red) in V1, the time segment highlighted in red for the M1 electrode was used.

### Multitaper analysis and non-negative matrix factorization extract states and their transitions across cortical layers and regions

To quantify the degree of coupling between state transitions at different recording sites, we developed a methodology to automatically detect state transitions at the level of individual channels (Methods). We then deployed this methodology to determine the degree to which transitions in different cortical sites are coupled. **Figure 3** is a flowchart of the initial analysis steps. The first step in the analysis is to compress the LFP recording into a low-sample-rate, low-dimensional matrix that accurately captures fluctuations in oscillatory activity. The right side of the figure presents an example five-minute window of data from one recording site to demonstrate the outcome of each step. Briefly, wideband data were filtered between 0.1 and 300 Hz to extract the LFP signal. (**Figure 3A**) LFP signals were converted to frequency domain using multitaper spectral analysis, (**Figure 3B**). Raw power spectra were then normalized such that the power contained in each frequency band was mapped onto a value between 0 (smallest observed power) and 1 (largest observed power) (**Figure 3C**). Non-negative matrix factorization (NMF) was used to further decompose the signal into a set of loadings and associated scores across time (**Figure 3D-E**).

**Figure 3.**
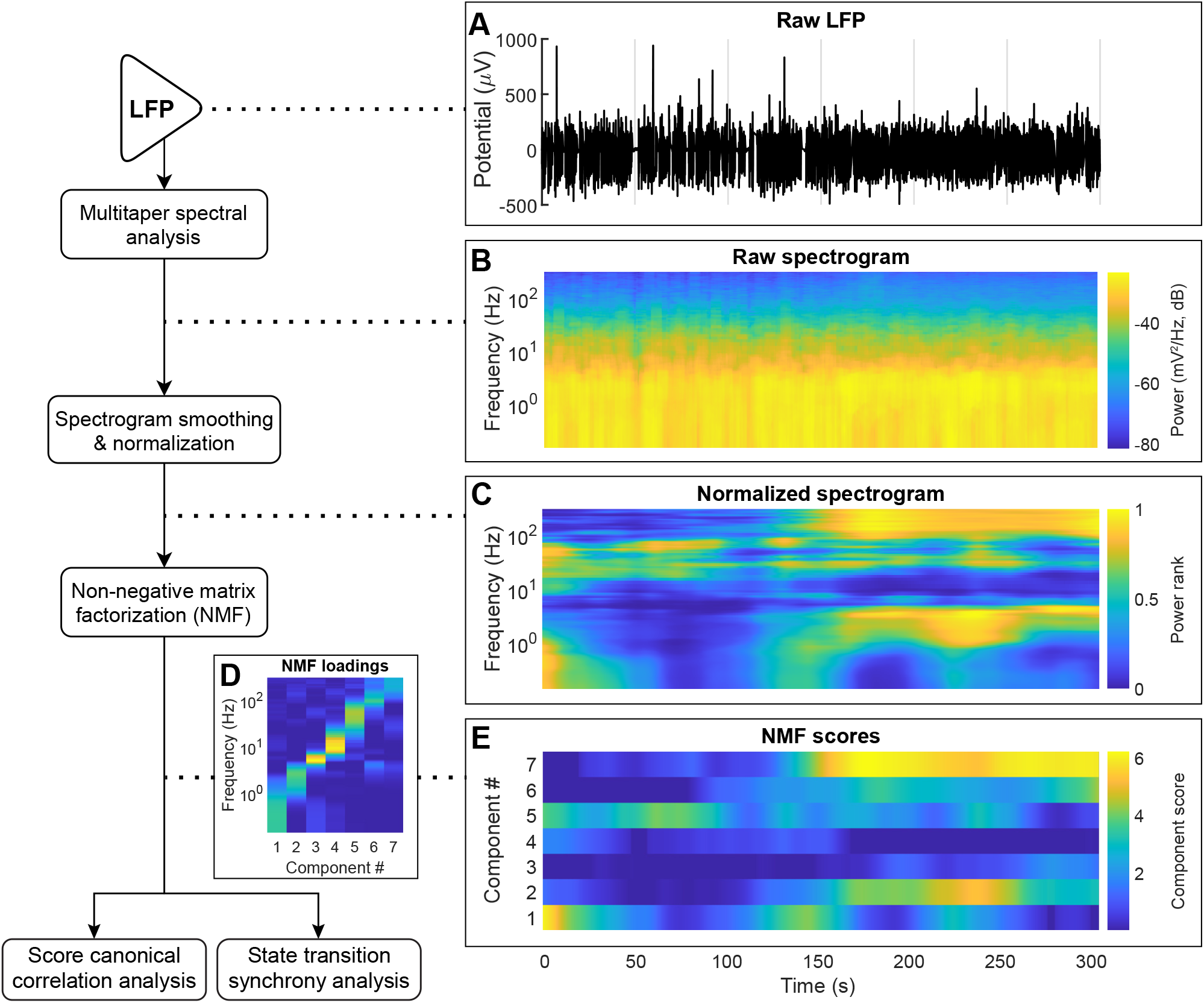
Schematic of LFP analysis, through NMF calculation. Left: Flowchart of analysis steps. Right: **A**. Five minutes of raw LFP signal centered around a state transition. **B**. Power spectrogram of LFP, computed using the multitaper method. **C**. The spectrogram from panel B after smoothing and rank-order normalization across time (Methods). **D**-**E** The loading (**D**) and score (**E**) matrices generated using NMF showing the spectral characteristics of each component and its relative contribution to the signal across time, respectively. The number of NMF components was optimized individually for each channel (Methods).

NMF can be thought of as a “soft” clustering algorithm. Previous work on state transitions under anesthesia (Hudson et al., 2014) and sleep (Gervasoni et al., 2004) used k-means clustering of the spectrograms to assign the state of the brain. Our first approach to state assignment used a similar strategy—the index of the NMF component with the highest score in each time window was defined as the state of the LFP at each recording site (Methods). This assumption was relaxed in subsequent stages of the analysis (see below). **Figure 4A** shows the score matrices for two different channels recorded simultaneously from two contacts along the same electrode in the motor cortex. The upper matrix is the same as **Figure 3E**, and the lower matrix was generated from data collected by a contact 140 um deeper inside the cortex. Notice that these matrices resemble one another but are not identical. **Figure 4B** shows state classifications for 18 channels of simultaneously recorded data: nine from an electrode in V1 and nine from an electrode in M1. Note again that some state transitions are observed around the same time in most of the electrodes. There are, however, many instances where state transition is observed in just a subset of the recording sites.

**Figure 4.**
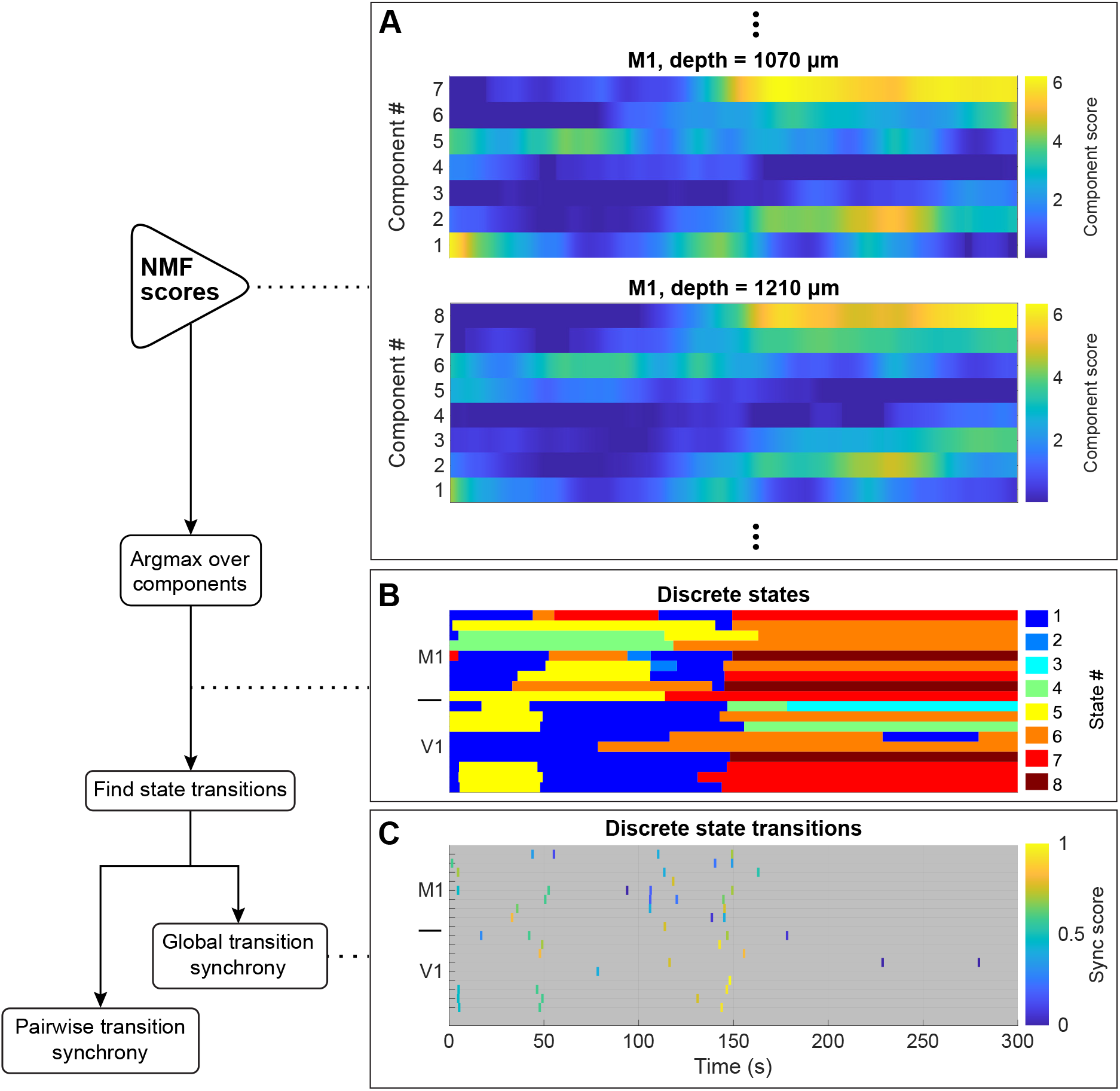
Schematic of NMF score analysis to define state transitions and synchrony. Left: Flowchart of analysis steps. Right: **A**. The NMF score matrix presented in Figure 3E (upper) and another NMF score matrix from simultaneously collected LFP from a neighboring channel (lower). Note that while nearby channels share similar characteristics across time, they are not identical. Also, the two channels have different optimal number of components, since NMF was performed and optimized (Methods) independently for each channel. **B**. State assignments across example time window from 18 simultaneously recorded signals: 9 signals from an M1 (top rows) electrode and 9 from V1 (bottom rows). State # indicates the NMF component with the highest score in each time window, after removing state segments that were both short and ambiguous due to small score fluctuations (Methods). **C**. Raster plot of all transition times from the channels presented in panel **B**. Transitions are colored according to their synchrony (sync score) with transitions in all other channels (Methods).

**Figure 5.**
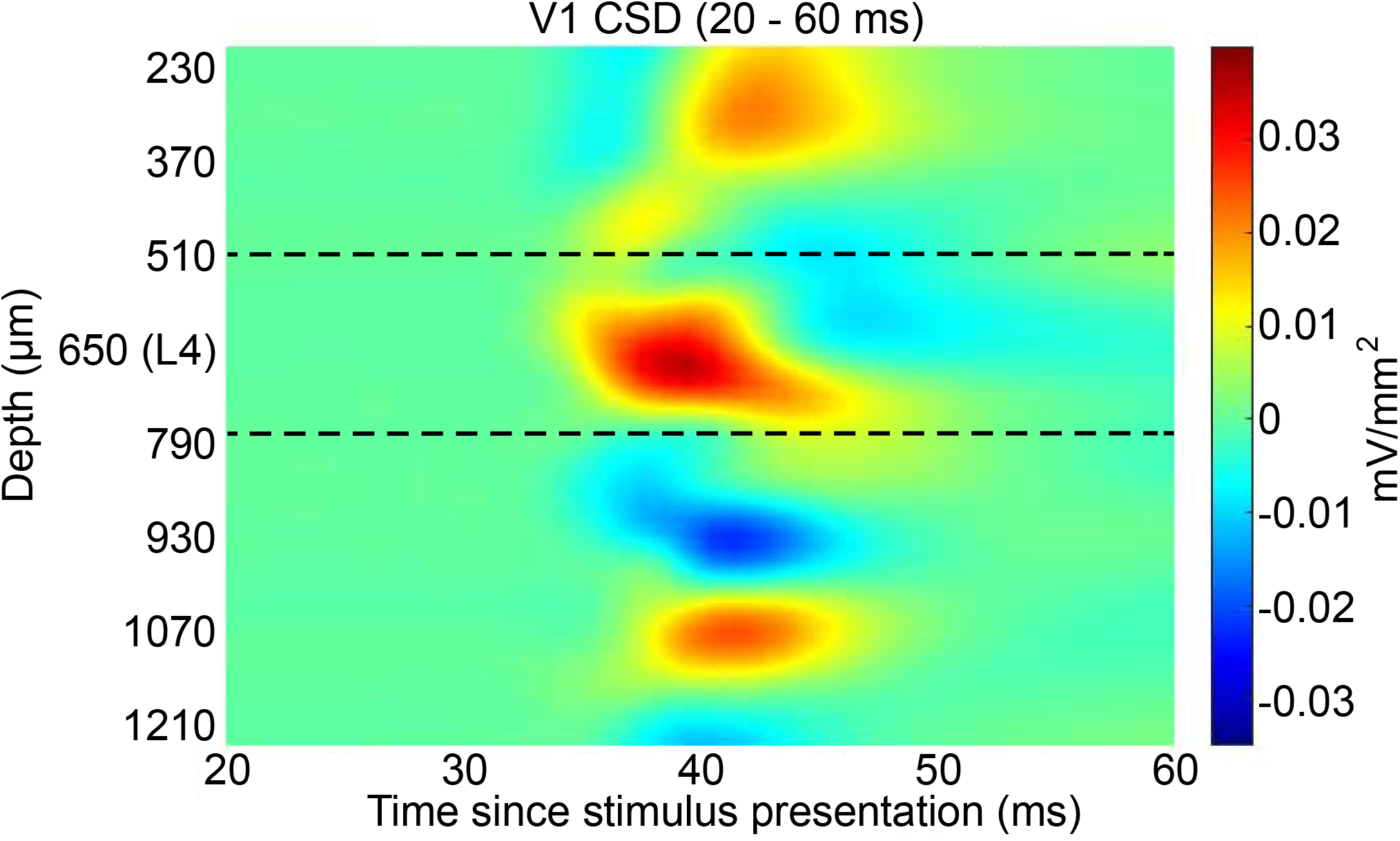
Current source density computed for a representative V1 recording. Evoked potential was elicited using a brief green LED flash (Methods). Dotted lines indicate the approximate boundaries of L4. Depth denotes estimated depth from the cortical surface.

**Figure 6.**
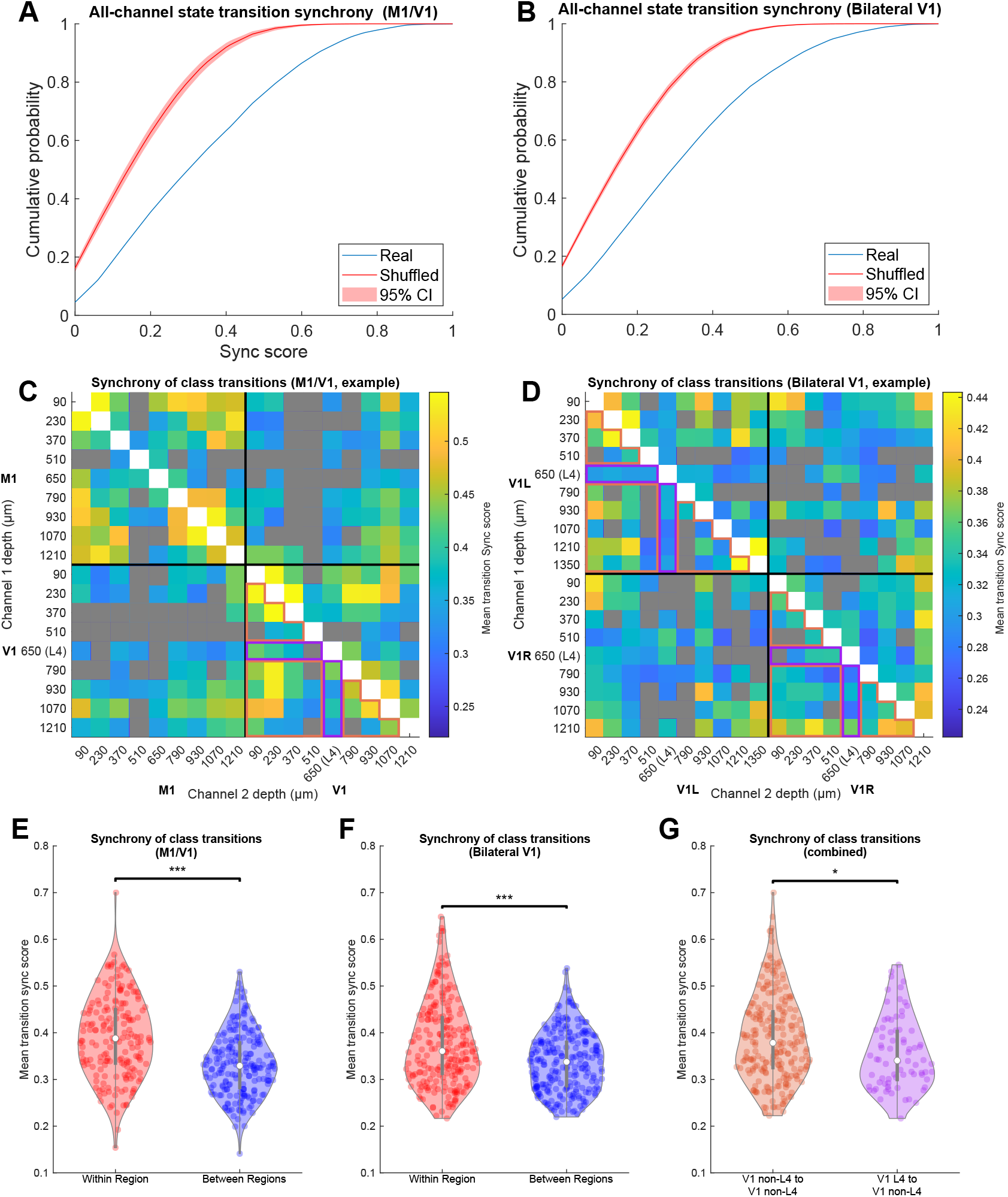
Transition synchrony between channels in the same anatomical region is higher than between channels in different regions. **A-B** The cumulative distribution of SPIKE-synchronization (synchrony) scores across all channels, in real recordings (blue) and the median ± 95% CI of 1000 shuffled recordings (red), for M1/V1 experiments (**A**) and bilateral V1 experiments (**B**). **C-D** Mean synchrony score across transitions for all channel pairs from a representative M1/V1 (**C**) and bilateral V1 (**D**) recording. Channel pairs whose synchrony scores were not significantly different from shuffled controls after Bonferroni correction are colored gray. **E-F** Channel pairs in which both channels are in the same region (red) have higher synchrony scores than those in which the channels are in different regions (blue) for M1/V1 (**E**, p = 1e-7, permutation test) and bilateral V1 (**F**, p = 2e-7, permutation test) recordings. **G**. Channel pairs in which one channel was within L4 and the other was not had lower synchrony scores than pairs in which neither channel was in L4 (p = 0.015, permutation test). Data included in these comparisons for the representative experiments are outlined in orange and purple, respectively, to highlight that only data from V1 electrodes were used.

**Figure 7.**
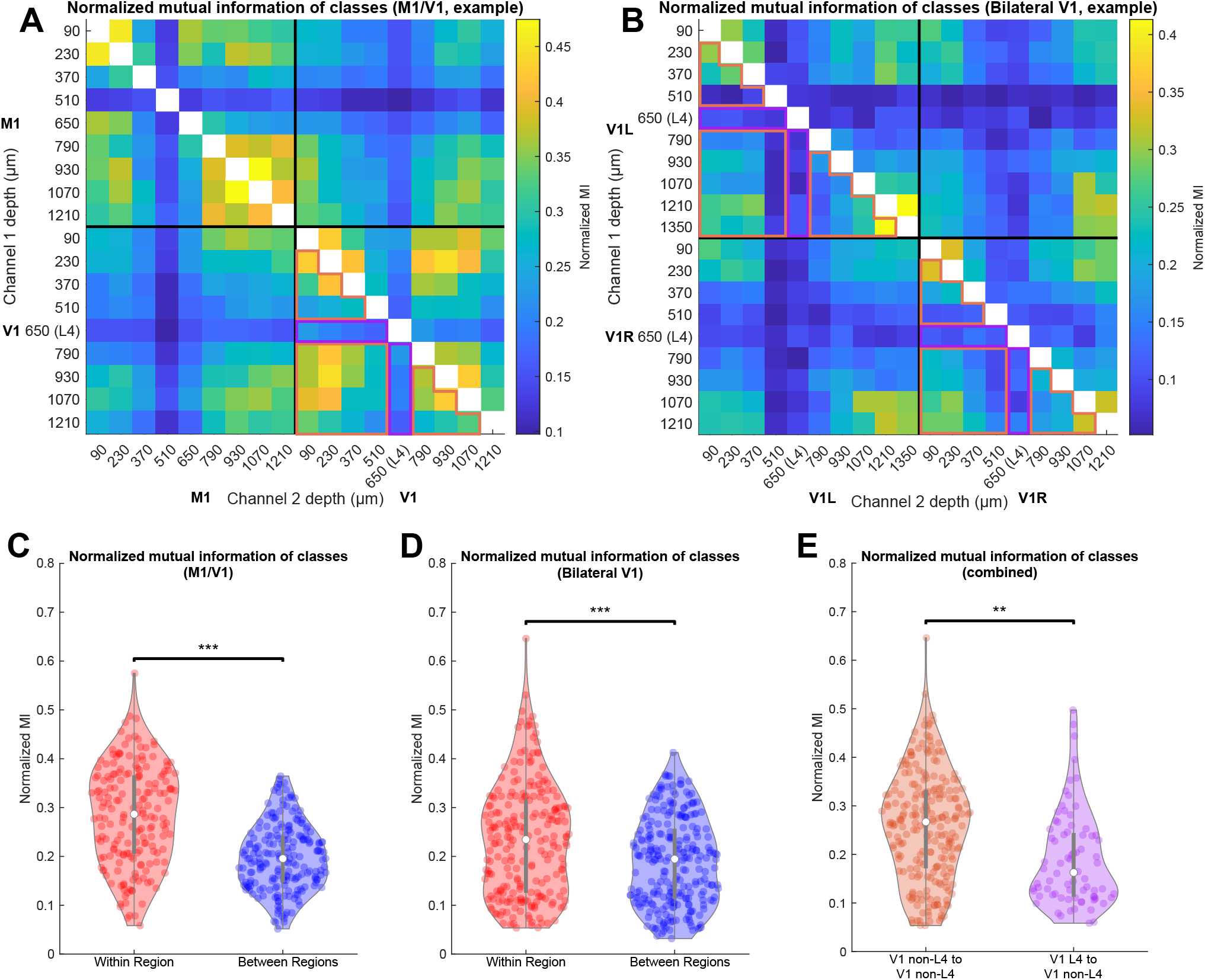
Normalized mutual information (MI) between channels in the same anatomical region is higher than between channels in different regions. **A-B** Normalized MI between state assignment vectors for all channel pairs from a representative M1/V1 (**A**) and bilateral V1 (**B**) recording. All normalized MI values are significantly different from shuffled controls after Bonferroni correction. **C-D** Channel pairs in which both channels are in the same region (red) have higher normalized MI than those in which the channels are in different regions (blue) for M1/V1 (**C**, p = 1e-7, permutation test) and bilateral V1 (**D**, p = 1e-7, permutation test) recordings. **E**. Channel pairs in which one channel was within L4 and the other was not had lower normalized MI than pairs in which neither channel was in L4 (p = 0.002, permutation test). Data included in these comparisons for the representative experiments are outlined in orange and purple, respectively, to highlight that only data from V1 electrodes were used.

**Figure 8.**
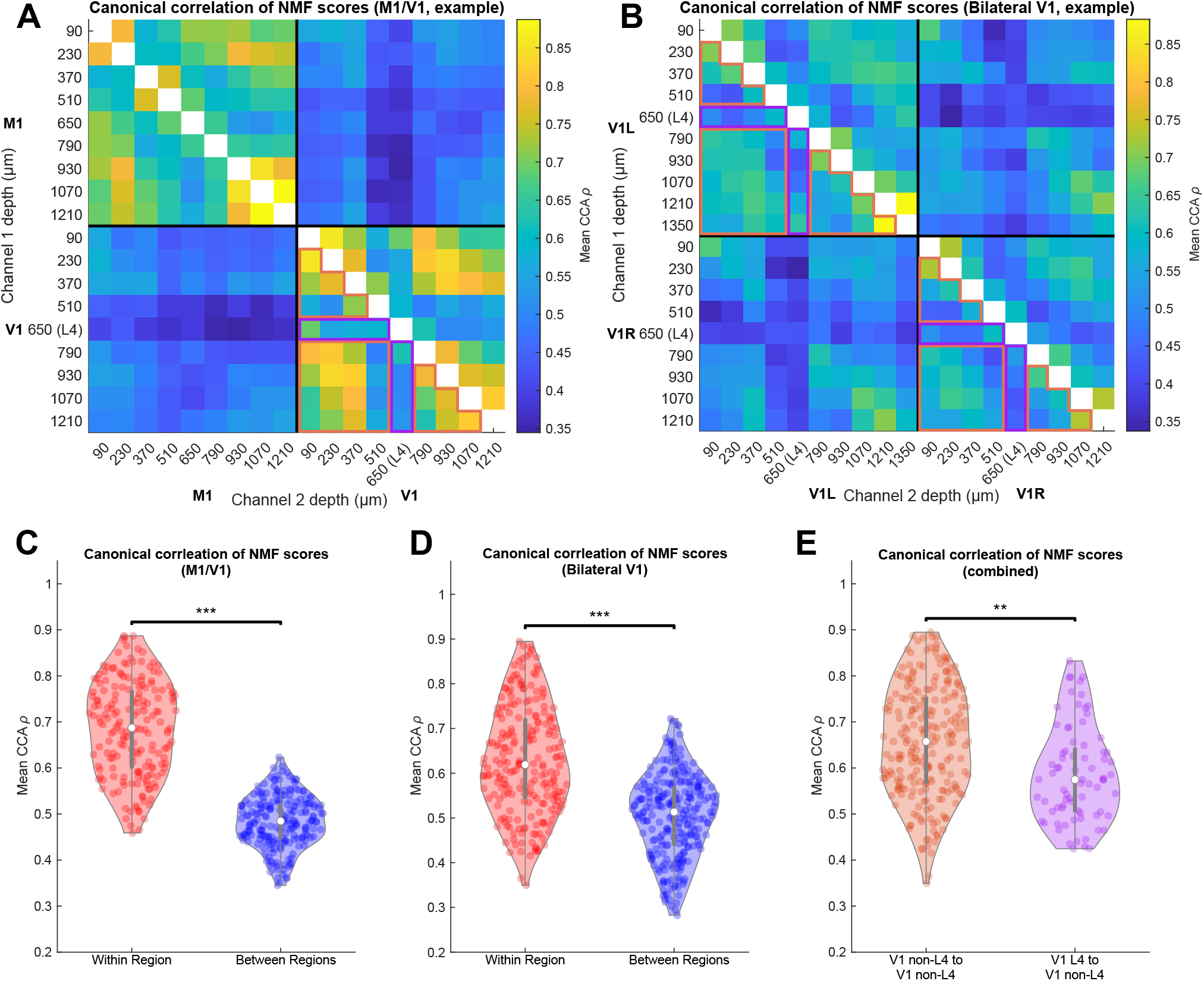
Canonical correlation analysis (CCA) reveals higher correspondence of overall activity between channels in the same anatomical region than between channels in different regions. **A-B** CCA measure on NMF scores for all channel pairs from representative M1/V1 (**A**) and bilateral V1 (**B**) recordings. **C-D** Channel pairs in which both channels are in the same region (red) have higher NMF score correspondence than those in which the channels are in different regions (blue) for M1/V1 (**C**, p = 1e-7, permutation test) and bilateral V1 (**D**, p = 1e-7, permutation test) recordings. **E**. Channel pairs in which one channel was within L4 and the other was not had lower NMF score correspondence than pairs in which neither channel was in L4 (p = 0.001, permutation test). Data included in these comparisons for the representative experiments are outlined in orange and purple, respectively, to highlight that only data from V1 electrodes were used.

One way to characterize the coupling between state transitions is to quantify the propensity of state transitions to occur simultaneously across different recording sites. Brain state transitions were defined as time points at which consecutive windows from the same channel have different brain state assignments (Methods). **Figure 4C** shows an example of this analysis. There are many transitions that appear in only one or very few channels, while others appear to be more global. **Figure 4C** is a raster plot of transitions. The color of each line shows the synchrony score of that transition with all other channels (Methods). Consistent with the observations in **Figure 2** and **4C**, the synchrony score reflects the fact that most state transitions are localized to a small subset of electrodes.

As we show below, coupling between state transitions depends on the cortical layer. Layer assignment in V1 was performed using current source density (CSD) analysis computed immediately following brief light stimulus (Methods). **Figure 5** shows a representative example of CSD in V1 showing the stereotypical pattern of response to visual stimuli. The first current sink occurs approximately 33 ms following stimulus presentation in L4. A short time after, additional sinks and sources appear above and below, revealing interlaminar communication. The channel where the initial sink occurred was defined as the center of L4. The dashed black lines in this figure mark the approximate boundaries of L4 based on the average thickness of this layer in rats and the spacing between channels (Einevoll et al., 2013; Quairiaux et al., 2011; Self et al., 2013). In the motor cortex, we did not estimate the location of cortical layers directly. Instead, we estimated the depth of each recording electrode relative to the cortical surface.

### State transitions in different cortical sites exhibit weak synchrony

We used three different analytical techniques to quantify the tendency of oscillatory states and the transitions between them to be coordinated across recording sites. Each technique relies on a different set of assumptions and was performed on a different feature of the data. First, we quantified the synchrony of transitions, as demonstrated in **Figure 4** (Methods). **Figure 6A-B** shows the cumulative distribution of synchrony scores (red curves) computed over all channel pairings and across all animals (M1/V1: 3 animals, 16–18 electrodes/animal, median of 99 transitions/electrodes/animal; bilateral V1: 4 animals, 15–19 electrodes/animal, median of 175.5 transitions/electrode/animal).

In order to compare the synchrony scores (**Figure 6A-B**) to those expected by chance, we generated shuffled datasets constrained to have the same state transition statistics. This was accomplished by simulating a Markov process defined by the state transition probability matrix derived from state assignments for each recording (Methods). This control preserves the statistics of each recording site, while destroying any coordination between them. The cumulative distributions of the synchrony scores obtained in these shuffled controls are shown in **Figure 6A-B** (blue curves; shading shows 95% confidence intervals computed over 1000 shuffled datasets). Both in the experiments involving M1 and V1 (**Figure 6A**) and in those involving bilateral V1s, we find that the synchrony score is consistently higher than expected by chance (p < 0.001, z-test based on means of shuffled datasets). Despite this large deviation from the null hypothesis, state transitions do not typically occur at the same time in different cortical sites (mean synchrony score ≈ 0.35 for both M1/V1 and bilateral V1 recordings). This implies that while state transitions observed across different cortical sites are not completely independent, coupling between channels is weak.

Data in **Figure 6A-B** aggregate the transition synchrony scores calculated between all channel pairs—both pairs of channels in the same cortical region and those located in different cortical sites. We hypothesized that, because most cortical connectivity is local, nearby electrodes would tend to have a higher propensity to change state at the same time. **Figure 6C-F** shows that state transitions are indeed more synchronous between electrodes within a cortical region than between regions. **Figure 6C-D** shows synchrony scores between all channel pairs in a representative pair of experiments: an M1/V1 experiment (**Figure 6C**) and a bilateral V1 experiment (**Figure 6D**). Pairs with scores that did not reach significance compared to the shuffled datasets, after Bonferroni correction for multiple comparisons, are shown in gray. Across all experiments, 57.0% of channel pairs from M1/V1 experiments and 80.2% of pairs from bilateral V1 experiments had significantly synchronous transitions at the corrected *p* < 0.05 level. The synchronization scores for all channel pairs from all experiments are quantified in **Figure 6E-F**, for M1/V1 and bilateral V1 experiments respectively. Both panels show the synchrony scores for within-region channel pairs (red) and between-region channel pairs (blue). In both types of recordings, within-region pairs had significantly larger synchrony scores than between-region pairs (*p =* 1e-7 for M1/V1 and *p =* 2e-7 for bilateral V1, compared to 10^7^ random permutations of the relevant channels (Methods)).

L4 is the thalamic input layer and has fewer horizontal connections than the supragranular or infragranular layers, which are rich in horizontal connections (Zilles and Palomero-Gallagher, 2017). To test whether layer organization affects transition synchrony, from each V1 recording (in which L4 was identified using CSD), we separated channel pairs in which one channel was in L4 from pairs in which neither channel was in L4. **Figure 6G** presents synchrony scores from all channel pairs from all experiments in which one channel was in L4 and the other was not (purple) and all channel pairs from all experiments in which neither channel was in L4 (orange). In **Figure 6E** and **F**, the specific channel pairs that were included in the “L4” and “non-L4” groups are outlined in purple and orange, respectively. We found that synchrony between channel pairs with one channel in L4 tended to be lower than between pairs in which neither channel was in L4 (*p =* 0.015, compared to 10^7^ random permutations of the relevant channels (Methods)). Therefore, transition times in channels from L4 tend to be relatively uncoupled from the specific timing of transitions in channels from other layers. This observation suggests that it is unlikely that thalamocortical input is the principal driver of state transitions in the cortex. If it were, one would expect that the thalamic input layer (L4) would transition in synchrony with the rest of the cortex. Therefore, these results imply different mechanisms, such as cortico-cortical interactions, are likely responsible for the timing of these spatially localized transitions.

Our final analysis using synchrony scores was performed to build upon these L4 results and determine whether the type of subcortical input to a cortical region has an influence on transition synchrony. It is typically assumed that switches of the oscillatory activity in the cortical LFP critically involve interactions with the thalamus (Contreras and Steriade, 1997; Herrera et al., 2016; Liu et al., 2015; Schiff, 2008; Steriade et al., 1993a, 1994). In light of this, one may expect two regions receiving similar thalamic input to exhibit greater synchrony of state transitions than two regions that interact with the thalamus in different ways. Therefore, we tested whether between-region comparisons for the bilateral V1 experiments had higher synchrony scores than the between-region comparisons for the M1/V1 experiments. Contrary to our hypothesis, we were not able to detect any increase in synchrony scores calculated between the bilateral V1s relative to M1/V1 experiments (*p =* 0.35, percentile bootstrap over channels (Methods)).

### Discrete states in different cortical sites have weak correspondence

Until this point, our analysis was based on transition synchrony, a measure that is sensitive to the timing of transitions but not the identities of the states. In what follows, we shift our focus away from the timing of state transitions and quantify the consistency of LFP-defined states at different sites. We accomplish this using normalized mutual information (MI), a measure of the amount of information obtained about one random variable by observing another random variable (Methods). In our case, these random variables are the time series of discrete states of two channels. High MI between these time series represents a large reduction in uncertainty about the state in channel *j* given the state in channel *i*. Two channels do not need to be in the same brain state to have high mutual information; indeed, since states are defined for each channel independently, there is no definition of different channels being in the “same” state. Rather, there must only be a consistent mapping from the states in one channel to those in the other. For example, if channel *i* is always in state A whenever channel *j* is in state D, one can predict the state of channel *i* from the state of channel *j*, and the MI between these channels would be high. As noted in the Methods, we normalized MI by the total entropy of the state distributions in the two channels over time in order to obtain a measure that was comparable across channels with different state distributions.

**Figure 7A-B** shows the normalized MI between all channel pairs in the same representative M1/V1 and bilateral V1 experiments as those in **Figure 6C-D**. 81.9% of channel pairs from M1/V1 experiments and 96.9% of pairs from bilateral V1 experiments had normalized MI that was significantly higher than for shuffled data, after Bonferroni correction for multiple comparisons (z-test based on distribution of shuffled data). The summary of normalized MI across all animals is shown in **Figure 7C-D**, for M1/V1 and bilateral V1 experiments respectively. In both types of recordings, within-region channel pairs had significantly higher normalized MI than between-region pairs (*p =* 1e-7 for M1/V1 and *p =* 1e-7 for bilateral V1, compared to 10^7^ random permutations of the relevant channels (Methods)). Note that, while for most channel pairs MI was higher than for a shuffled dataset, the amount of information about the state of one channel contained in the state of another was small. Normalized mutual information varies between 0 and 1, where 1 denotes that the two channels carry identical information. Yet, even in a pair of channels within a single cortical region, the mean MI is about 0.3. One way to interpret this statistic (Methods) is that no more than 15% of the combined information carried by the states of any two channels is redundant. Thus, most of the information about the state of one channel cannot be extracted from observing the state of a nearby channel in the cortex.

As with transition synchrony, we did not detect a higher mean normalized MI in left/right V1 channel pairs compared to M1/V1 channel pairs (*p* = 0.70, percentile bootstrap over channels (Methods)). Additionally, as with the transition synchrony analysis, pairs including a channel in L4 did have lower normalized MI than pairs where neither channel was in L4 (*p* = 0.002, compared to 10^7^random permutations of the relevant channels (Methods)). These results show not only that channels from the same brain region are more likely to undergo transitions at the same time, but also that the broader structure of these state assignments across the entire recording is more similar in channels from the same region. Furthermore, the conclusions regarding the differences between L4 and other cortical layers are consistent between synchrony and mutual information analyses.

### Full compressed spectrograms of different sites have moderate correspondence, depending on distance and cortical layer

In the previous analyses, to generate a single-value description of activity across time, we defined brain state as the NMF loading with the highest score in each time window. This method was convenient for comparing synchrony of transitions and mutual information of state sequences. Parcellation of the LFP signals into discrete states is also supported by previous work (Hudson et al., 2014) However, reducing the LFP to a single value eliminates much of the information in the original signal. In order to incorporate more of this information, rather than collapsing the LFP signal to a single value, we used the vector of NMF scores for the LFP in each temporal window directly. Each score vector, once multiplied through by the appropriate loading matrix (Methods and **Figure 3**), yields a good approximation of the actual spectrum of the LFP in that time window. To test for correlated fluctuations in the spectral features of LFPs at different cortical sites, we applied canonical correlation analysis (CCA) to the pair of score matrices derived from each pair of channels. High canonical correlation indicates a close linear relationship between two sets of variables. The mean of the vector *ρ* of canonical correlations between all pairs of canonical variables was calculated to give a measure of overall state similarity that is invariant to invertible linear transformations of each channel’s state space. This method of taking the average across *ρ* is explained further in Alpert and Peterson (1972). **Figure 8A-B** shows the CCA similarity measure for all channel pairs from the same representative M1/V1 and bilateral V1 experiments that have been shown previously. All channel pairs from both M1/V1 and bilateral V1 experiments had significantly higher CCA similarities than for shuffled data, after Bonferroni correction for multiple comparisons (z-test based on distribution of shuffled data). The summary of CCA similarity across all animals is shown in **Figure 8C-D**. These results are very similar to those for transition synchrony and normalized MI and show that in both types of recordings, within-region channel pairs had significantly higher CCA similarities than between-region pairs (*p =* 1e-7 for M1/V1 and *p =* 1e-7 for bilateral V1, compared to 10^7^ random permutations of the relevant channels (Methods)). Furthermore, as with the previous measures, channel pairs including a channel in L4 had lower CCA similarities than pairs in which neither channel was in L4 (*p =* 0.001, compared to 10^7^ random permutations of the relevant channels (Methods)). We did not detect a higher mean CCA similarity in left/right V1 channel pairs compared to M1/V1 channel pairs (*p =* 0.12, percentile bootstrap over channels (Methods)).

### Global brain state is low-dimensional, despite weak pairwise interactions

All results shown up until this point were calculated on pairs of channels for which state assignments were computed independently. What we have shown is that channels within the same cortical region tend to be more similar in their activity patterns and state transition times than channels from different cortical regions. However, close inspection of the results shows that, even for the channel pairs within the same cortical region, only about one third of the information contained within the discrete state sequences is shared between channels (**Figure 7C**). For channel pairs from different cortical regions, the amount of mutual information in state sequences is even lower. This weak coupling between channels could imply that spatially restricted regions of the brain act independently of one another and there is no discernable global state of the brain at any given time. Alternatively, it is possible that this weak coupling between channels, *en masse*, gives rise to a complex, global state of activity that is differently expressed in the oscillation patterns of spatially restricted regions of cortex. In this final analysis, we sought to directly distinguish these possibilities by characterizing the global brain state. In a key distinction from the previous work, rather than defining the global state on the basis of the concatenated spectra from all recordings, we attempted to identify global macroscopic dynamics from the simplified dynamics observed at each recording site. This was accomplished by first concatenating the NMF score vectors from all simultaneously recorded channels at each timepoint into a single vector that encodes the joint state of all channels. The resulting full matrix of joint states over time was then subjected to principal component analysis (PCA).

We found that all but one recording required 10 or fewer components to account for 80% of the variance in the concatenated NMF score matrices, which ranged in dimensionality from 91 to 136. The recording that required greater than 10 required 15 components to reach the same threshold. This is far outside the 95% confidence interval of expected cumulative explained variance, computed on Markov-shuffled controls which ignore weak pairwise correlations between fluctuations in different channels (**Figure 9A, D**). These results demonstrate that widespread weak coupling is sufficient to give rise to a highly correlated global state. **Figures 9B** and **E** show the loadings onto channels and frequencies (mapped back from corresponding NMF loadings) for the top two principal components of a representative M1/V1 and bilateral V1 recording, respectively. These data offer qualitative evidence that the global state is differentially reflected in different regions and layers of the cortex. For example, the loadings of the second principal component (PC2) of the M1/V1 recording in **Figure 9B** show that, while there is high power in the higher frequencies for the V1 channels, the same is not true in the M1 channels. In contrast,

**Figure 9.**
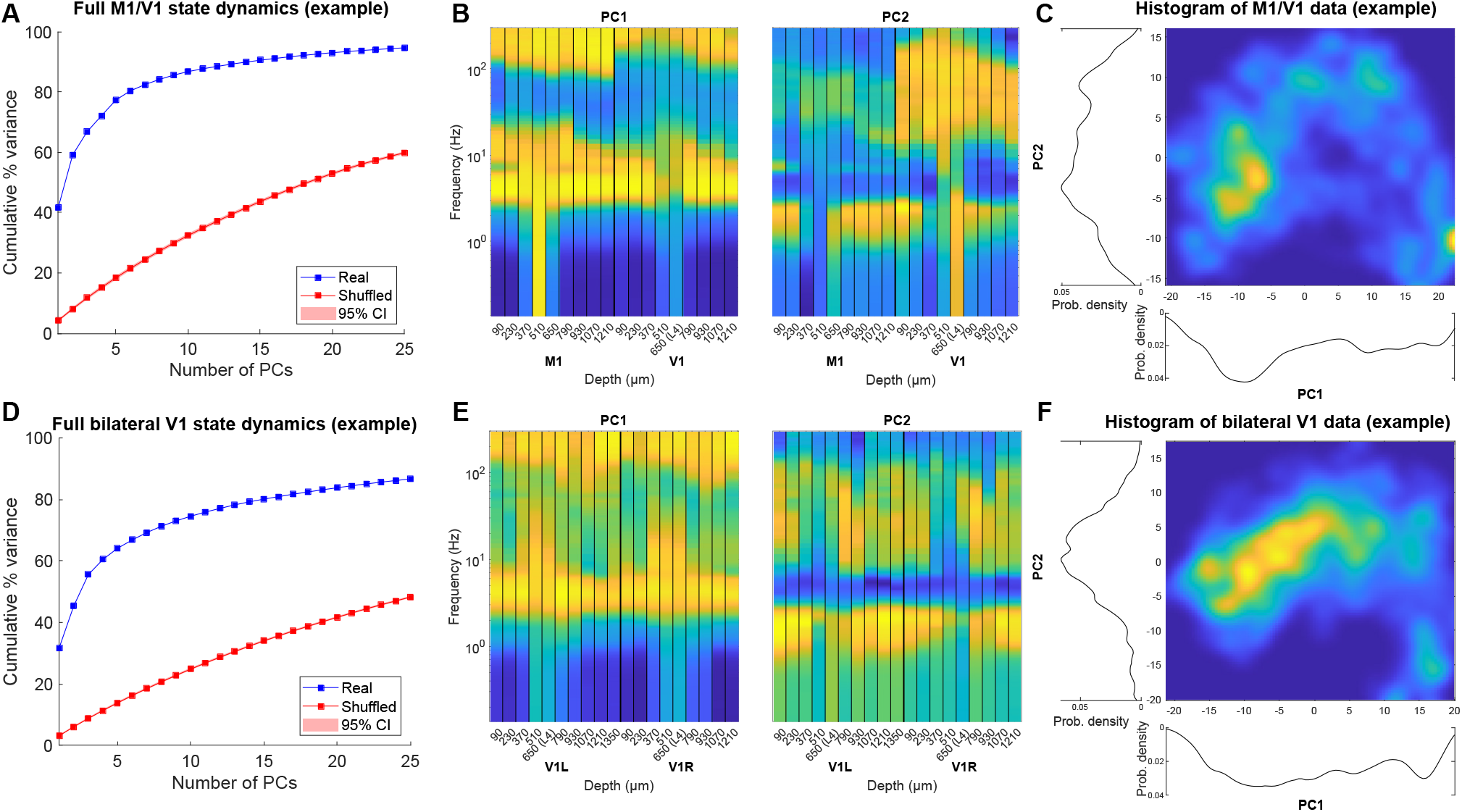
Weakly correlated fluctuations in different cortical sites give rise to highly correlated cortical states. NMF scores from all recorded channels were concatenated into a single state vector (median dimension across recordings = 106) and subjected to PCA. Fraction of total variance as a function of number of PCs is shown in **A** and **D** for M1/V1 and bilateral V1 example recordings respectively (blue). Shuffled surrogates (Methods) were subjected to the same analysis (red). **B** and **E** show loadings of the top 2 principal components, mapped back from each channel’s NMF components to frequencies, for the two representative recordings. This projection reveals consistent differences between M1 and V1 (**B**) but is relatively consistent across bilateral V1s (**E**). In both instances, Layer 4 is distinct from supra- and infragranular layers. **C** and **F** show histograms of the data projected onto the top two PCs for the representative M1/V1 (**C**) and bilateral V1 (**F**) recordings. In both instances, the distribution of data is multimodal, suggesting the presence of discrete global cortical states.

**Figure 9E** shows that the loadings of PC1 of the bilateral V1 recording onto all channels of both electrodes are fairly uniform, except for in channels near L4 where there is higher power in the lowest frequency bands. **Figures 9C** and **F** show histograms of all samples from these representative recordings projected onto the first two principal components. Although more than two dimensions would be necessary to fully visualize the landscape of the global dynamics, even in this limited projection, a clustered pattern is visible, similar to previous results (Hudson et al., 2014). These data suggest that global brain states comprise regionally distinct oscillation patterns that are weakly coupled with one another. Remarkably, these results show that discrete transitions between global cortical states (Ballesteros et al., 2020; Hudson et al., 2014; Patel et al., 2020) under a fixed anesthetic concentration arise from the multitude of weakly coupled local fluctuations.

## Discussion

Here we set out to determine how abrupt transitions between global thalamocortical states arise at a fixed anesthetic concentration. Using several complementary analysis methods, we demonstrate that correlated fluctuations in the oscillatory behavior observed at different cortical sites are widespread, but that each pairwise interaction is weak. Thus, for instance, the ability to infer the current state of one channel by observing the state of a nearby channel in the cortex is limited. Remarkably, we provide evidence that abrupt transitions between discrete macroscopic cortical activity patterns (Ballesteros et al., 2020; Chander et al., 2014; Hudson et al., 2014; Ishizawa et al., 2016; Lee et al., 2020; Patel et al., 2020) emerge naturally from the multitude of these quasi-independent local fluctuations. We also demonstrate that the strength of the interactions between recording sites depends on the inter-electrode distance and on the cortical layer. Specifically, we find that fluctuations in L4, the thalamic input layer, tend to be less congruent with those in other layers. Altogether, these results argue that abrupt global state transitions are not imposed on the thalamocortical networks by changes in the activity of broadly projecting modulatory arousal systems, but rather are strongly influenced by the local cortico-cortical interactions.

It has been conjectured that structured transitions between discrete states constrain the space of possible brain activity patterns and thereby allow the brain to efficiently recover its normal waking state after a dramatic perturbation (Hudson et al., 2014). The idea that, in order to recover from a perturbation, the space of possible activity states must be constrained by stabilization of a few discrete activity patterns is not specific to recovery from anesthesia *per se*. For instance, pharmacologically provoked recovery of consciousness in the setting of brain injury is also characterized by abrupt transitions between quasi-stable activity patterns (Victor et al., 2011). Sleep is also well known to consist of discrete activity patterns (e.g., Gervasoni et al., 2004). Thus, it appears that abrupt state transitions among discrete activity states accompany recovery of normal consciousness in a variety of settings.

It is thus of great interest to determine how such state transitions arise and how they are coordinated across thalamocortical networks. Here, in keeping with previous work (e.g., Gervasoni, 2004; Hudson et al., 2014), we defined the state of each local recording site on the basis of the power spectrum of the LFP. Since we focused on state transitions observed in the anesthetized brain, most fluctuations occurred in the slow oscillations (< 1 Hz) (Steriade et al., 1993b), delta oscillations (1-4 Hz), and the spindle range of 8-14 Hz (Purpura, 1968). Multiple distinct neurophysiological mechanisms contribute to the generation and coordination of the various brain oscillations observed in the anesthetized brain. Slow oscillations, for instance are thought to be primarily generated through local synaptic mechanisms in the cortex (Sanchez-Vives and McCormick, 2000; Steriade et al., 1993b). Thalamocortical and thalamic reticular neurons reflect these slow oscillations and are phase locked to them (Steriade et al., 1993b). However, the fact that slow oscillations are abolished in the thalamus of decorticated animals (Timofeev and Steriade, 1996) but are observed in the cortex of athalamic animals (Steriade et al., 1993b) strongly argues for the cortical origin of slow oscillations. Cortico-cortical interactions are thought to underlie not just the generation of slow waves, but also the synchronization of these waves across the cortex. Pharmacologic and surgical lesions of intra-cortical connections disrupt the synchrony of slow waves (Amzica and Steriade, 1995).

The observation that slow oscillations are coordinated primarily through cortico-cortical interactions is consistent with our results. Many of the state transitions under isoflurane involve fluctuations in the power of slow oscillations. Using three distinct analysis methods, we consistently find that state fluctuations in L4 are relatively dissimilar to those observed in the infra- and supragranular layers. L4 neurons are most directly affected by spatially localized inputs from the thalamus, whereas supra- and infra-granular neurons are primarily driven by cortico-cortical connections and matrix projections from the thalamus (Jones, 2001). While anesthetics suppress both core and matrix thalamocortical inputs, their dominant effect is specifically suppressing cortico-cortical connectivity (Raz et al., 2014). It is thus likely that the local nature of state transitions in the slow oscillation range is a consequence of both weakened thalamocortical and cortico-cortical interactions in the anesthetized brain.

Transitions between slow (< 4Hz) and faster EEG oscillations, occasionally observed even in the anesthetized brain (e.g., **Figure 2**) are thought to arise as a result of the interaction of the thalamo-cortical networks with neuromodulatory projections from cholinergic neurons in the brainstem and basal forebrain (Steriade, 2004). Noradrenergic neurons (Vazey and Aston-Jones, 2014) and other brain stem and basal forebrain nuclei also contribute to the modulation of the oscillations exhibited by the thalamocortical networks (Jones, 2003). Activity within the various arousal promoting nuclei is coordinated by a group of medullary neurons, activation of which can trigger prompt awakening from deep states of anesthesia (Gao et al., 2019). In the anesthetized brain, fluctuations in the firing rate of these medullary neurons co-varies with the fluctuations in the spectral characteristics of the cortical LFP (Gao et al., 2019). Thus, it is possible that the spontaneous fluctuations of the LFP characteristics between the slower and faster oscillations are in part mediated by fluctuations in the activity of the nuclei that modulate the thalamocortical networks. However, most arousal nuclei have broad projections to the thalamus and the cortex (Jones, 2003). Thus, if the fluctuations in the state of the LFP were entirely driven by the fluctuations in the activity of the modulatory projections, one would expect that the state of the LFP would fluctuate coherently across the cortex. Instead, we observe that fluctuations in state of the LFP are only weakly coupled between different cortical sites. This implies that the influence of the modulatory nuclei on the power of specific cortical oscillations, within the physiological range, is not absolute. Rather, activation of the modulatory systems likely biases the cortex towards a particular oscillatory state. The overall pattern of activity at each cortical site, however, is strongly influenced by interactions within the thalamocortical networks.

The experiments performed here cannot directly address the cellular and synaptic mechanisms that give rise to local state transitions and their coordination across the cortex. They do, however, offer clear insights into network mechanisms of global state transitions. Here, rather than attempting to simplify the dynamics of the global signals directly (Hudson et al., 2014), we embedded the dynamics of the local signals into a low-dimensional space. This analysis revealed only weak interactions between local signals. Remarkably, assembling just the low-dimensional projections of the local signals into a state vector recapitulated the low-dimensional dynamics and discrete global cortical states. Thus, we show that the global states and abrupt transitions between them arise because of weak coupling between local state fluctuations.

We are not the first to note that weak coupling among local fluctuations can give rise to coherent macroscopic states. In the retina, weak correlations in spike timing co-exist with a conspicuously high probability of certain large ensembles of neurons firing in synchrony (Schneidman et al., 2006). It may seem that a network with weakly correlated nodes can be well approximated by a collection of completely independent nodes, but this is not the case. Weakly coupled elements can yield highly correlated macroscopic states if the weak interactions are prevalent enough throughout the network. Indeed, we find that while the correlations between different cortical sites were weak, they were present and statistically significant for most electrode pairs.

The emergence of highly correlated global states from weak pairwise interactions has been investigated extensively in statistical mechanics using Ising models. It has been shown that an Ising model is mathematically equivalent to a maximum entropy models of the statistics of neural firing that are constrained only by the experimentally observed firing probabilities of individual neurons and their pairwise correlations (Schneidman et al., 2006; Tkačik et al., 2006). The maximum entropy approach has proved successful in diverse systems (Ohiorhenuan et al., 2010; Tang et al., 2008; Tkačik et al., 2014; Yu et al., 2008). Although Ising models have traditionally been applied to binary state spaces, such as the presence or absence of an action potential within a small time window, the maximum entropy approach can be generalized to continuous variables (Bialek et al., 2012), such as local fields. In this work, we did not explicitly attempt to construct a maximum entropy model of local field fluctuations, as we are recording only a tiny fraction of all cortical signals. Future work may sample of local field fluctuations more densely to determine whether an Ising-type model suffices to explain the fluctuations of the global state of the brain under anesthesia, or whether other mechanisms in addition to pairwise interactions are needed (Ohiorhenuan et al., 2010; Tang et al., 2008). Regardless of the specific details of such a model, however, we directly demonstrate that widespread weak correlations in local field fluctuations give rise to coherent global cortical states. This conclusion is strongly supported by the observations that locally defined cortical states yield highly correlated global behavior despite weak pairwise interactions, whereas the shuffled controls do not.

There are multiple parallels between our characterization of state transitions in the anesthetized brain and those observed during slow wave sleep (NREM). While sleep and anesthesia are clearly distinct phenomena, the neurophysiological mechanisms that give rise to oscillations in the thalamocortical circuitry under anesthesia and during natural sleep share some essential similarities (Steriade et al., 1993b; Steriade and Amzica, 1998). Many diverse anesthetics promote activity in the sleep active subcortical nuclei and suppress activity in the wake active ones (Jiang-Xie et al., 2019; Moore et al., 2012; Nelson et al., 2002; Zhang et al., 2015). Furthermore, both sleep and anesthesia consist of several discrete states, each characterized by a distinct pattern of oscillations in the cortex and thalamus (Saper et al., 2010). Based on the original recordings at the microscopic level of single isolated neurons or, alternatively, on the macroscopic level using EEG, it has long been hypothesized that sleep stages are brain-wide phenomena and that the neurophysiological mechanisms that give rise to sleep stage switching specifically prevent multiple sleep stages or sleep and wakefulness from coexisting at the same time in different brain regions (Lu et al., 2006; Saper et al., 2010). Interestingly, at the mesoscopic level of neuronal populations and local fields, sleep state transitions, much like in this work, can be local (Nir et al., 2011; Poulet and Petersen, 2008; Vyazovskiy et al., 2011). Furthermore, it has been suggested that antecedent neuronal activity driven by a specific task can increase the propensity of a population of cortical neurons to exhibit local sleep-like slow oscillations (Huber et al., 2004), implying that transitions between different oscillatory modes are strongly influenced by local synaptic interactions. The degree of synchrony between cortical locations across naturally observed state transitions, such as those between different sleep stages or between sleep and wake, has not been directly quantified in a systematic fashion. Because sleep is strongly influenced by both homeostatic and circadian influences, it will be challenging to disentangle these global influences from the local interactions between different sites in the cortex. However, analysis of cortical state transitions in the brain anesthetized with a fixed anesthetic concentration is free from these complications. This analysis shows that the apparently global coordinated shifts in cortical activity arise naturally out of weakly interacting local state switches.

## Data and Software Availability

Datasets are available upon reasonable request. Code for time-frequency analysis of LFP and interaction measure calculation is publicly available at https://github.com/ProektLab/spec-state-trans and other public repositories linked from the README.

## Acknowledgements

The authors thank Andrew Hudson for his implementation of the multitaper method, as well as Adeeti Aggarwal, Max Kelz, Andrew McKinstry-Wu, and Andi Wasilczuk for helpful conversations. This work was supported by National Institutes of Health R01 grants no. 5R01GM124023 and 5R01NS113366. B.P.S. was also supported by an NIH National Research Service Award (F31) (grant no. 1F31NS118808-01A1).

## Works Cited

Alpert MI, Peterson RA. 1972. On the Interpretation of Canonical Analysis. Journal of Marketing Research 9:7.

Amzica F. 2009. Basic physiology of burst-suppression. Epilepsia 50:38–39. doi:10.1111/j.1528-1167.2009.02345.x

Amzica F, Steriade M. 1995. Disconnection of intracortical synaptic linkages disrupts synchronization of a slow oscillation. J Neurosci 15:4658–4677. doi:10.1523/JNEUROSCI.15-06-04658.1995

Ballesteros JJ, Briscoe JB, Ishizawa Y. 2020. Neural signatures of α2-Adrenergic agonist-induced unconsciousness and awakening by antagonist. eLife 9:e57670. doi:10.7554/eLife.57670

Bialek W, Cavagna A, Giardina I, Mora T, Silvestri E, Viale M, Walczak AM. 2012. Statistical mechanics for natural flocks of birds. PNAS 109:4786–4791. doi:10.1073/pnas.1118633109

Brown EN, Lydic R, Schiff ND. 2010. General Anesthesia, Sleep, and Coma. New England Journal of Medicine 363:2638–2650. doi:10.1056/NEJMra0808281

Canavier CC, Baxter DA, Clark JW, Byrne JH. 1993. Nonlinear dynamics in a model neuron provide a novel mechanism for transient synaptic inputs to produce long-term alterations of postsynaptic activity. J Neurophysiol 69:2252–2257. doi:10.1152/jn.1993.69.6.2252

Chander D, García PS, MacColl JN, Illing S, Sleigh JW. 2014. Electroencephalographic Variation during End Maintenance and Emergence from Surgical Anesthesia. PLoS ONE 9:e106291. doi:10.1371/journal.pone.0106291

Civillico EF, Contreras D. 2012. Spatiotemporal properties of sensory responses in vivo are strongly dependent on network context. Frontiers in Systems Neuroscience 6:1–20. doi:10.3389/fnsys.2012.00025

Contreras D, Steriade M. 1997. State-dependent fluctuations of low-frequency rhythms in corticothalamic networks. Neuroscience 76:25–38. doi:10.1016/s0306-4522(96)00392-2

Destexhe A, Contreras D, Sejnowski TJ, Steriade M. 1994. Modeling the control of reticular thalamic oscillations by neuromodulators. Neuroreport 5:2217–2220. doi:10.1097/00001756-199411000-00003

Einevoll GT, Kayser C, Logothetis NK, Panzeri S. 2013. Modelling and analysis of local field potentials for studying the function of cortical circuits. Nature Reviews Neuroscience 14:770–785.

Ermentrout B. 1998. Neural networks as spatio-temporal pattern-forming systems. Rep Prog Phys 61:353–430. doi:10.1088/0034-4885/61/4/002

Fisher RS, Engel JJ. 2010. Definition of the postictal state: When does it start and end? Epilepsy & Behavior 19:100–104. doi:10.1016/j.yebeh.2010.06.038

Friedman EB, Sun Y, Moore JT, Hung H-T, Meng QC, Perera P, Joiner WJ, Thomas SA, Eckenhoff RG, Sehgal A, Kelz MB. 2010. A Conserved Behavioral State Barrier Impedes Transitions between Anesthetic-Induced Unconsciousness and Wakefulness: Evidence for Neural Inertia. PLoS One 5:e11903. doi:10.1371/journal.pone.0011903

Gao S, Proekt A, Renier N, Calderon DP, Pfaff DW. 2019. Activating an anterior nucleus gigantocellularis subpopulation triggers emergence from pharmacologically-induced coma in rodents. Nat Commun 10:2897. doi:10.1038/s41467-019-10797-7

Gervasoni D, Lin S-C, Ribeiro S, Soares ES, Pantoja J, Nicolelis MAL. 2004. Global Forebrain Dynamics Predict Rat Behavioral States and Their Transitions. J Neurosci 24:11137–11147. doi:10.1523/JNEUROSCI.3524-04.2004

Herrera CG, Cadavieco MC, Jego S, Ponomarenko A, Korotkova T, Adamantidis A. 2016. Hypothalamic feedforward inhibition of thalamocortical network controls arousal and consciousness. Nature Neuroscience 19:290–298. doi:10.1038/nn.4209

Hodgkin AL, Huxley AF. 1952. A quantitative description of membrane current and its application to conduction and excitation in nerve. J Physiol 117:500–544.

Huber R, Felice Ghilardi M, Massimini M, Tononi G. 2004. Local sleep and learning. Nature 430:78–81. doi:10.1038/nature02663

Hudson AE, Calderon DP, Pfaff DW, Proekt A. 2014. Recovery of consciousness is mediated by a network of discrete metastable activity states. Proceedings of the National Academy of Sciences 111:9283–9288. doi:10.1073/pnas.1408296111

Ishizawa Y, Ahmed OJ, Patel SR, Gale JT, Sierra-Mercado D, Brown EN, Eskandar EN. 2016. Dynamics of Propofol-Induced Loss of Consciousness Across Primate Neocortex. Journal of Neuroscience 36:7718–7726. doi:10.1523/JNEUROSCI.4577-15.2016

Izhikevich EM. 2007. Dynamical systems in neuroscience: the geometry of excitability and bursting, Computational neuroscience. Cambridge, Mass: MIT Press.

Jiang-Xie L-F, Yin L, Zhao S, Prevosto V, Han B-X, Dzirasa K, Wang F. 2019. A Common Neuroendocrine Substrate for Diverse General Anesthetics and Sleep. Neuron 102:1053-1065.e4. doi:10.1016/j.neuron.2019.03.033

Joiner WJ, Friedman EB, Hung H-T, Koh K, Sowcik M, Sehgal A, Kelz MB. 2013. Genetic and Anatomical Basis of the Barrier Separating Wakefulness and Anesthetic-Induced Unresponsiveness. PLoS Genet 9:e1003605. doi:10.1371/journal.pgen.1003605

Jones BE. 2003. Arousal systems. Front Biosci 8:s438–451. doi:10.2741/1074

Jones EG. 2001. The thalamic matrix and thalamocortical synchrony. Trends Neurosci 24:595–601. doi:10.1016/s0166-2236(00)01922-6

Kelz MB, Sun Y, Chen J, Cheng Meng Q, Moore JT, Veasey SC, Dixon S, Thornton M, Funato H, Yanagisawa M. 2008. An essential role for orexins in emergence from general anesthesia. Proc Natl Acad Sci U S A 105:1309–1314. doi:10.1073/pnas.0707146105

Kreuz T, Bozanic N, Mulansky M. 2015. SPIKE-Synchronization: a parameter-free and time-resolved coincidence detector with an intuitive multivariate extension. BMC Neurosci 16:P170, 1471-2202-16-S1-P170. doi:10.1186/1471-2202-16-S1-P170

Lee DD, Seung HS. 1999. Learning the parts of objects by non-negative matrix factorization. Nature 401:788–791. doi:10.1038/44565

Lee H, Wang S, Hudetz AG. 2020. State-Dependent Cortical Unit Activity Reflects Dynamic Brain State Transitions in Anesthesia. J Neurosci 40:9440–9454. doi:10.1523/JNEUROSCI.0601-20.2020

Liu J, Lee HJ, Weitz AJ, Fang Z, Lin P, Choy M, Fisher R, Pinskiy V, Tolpygo A, Mitra P, Schiff N, Lee JH. 2015. Frequency-selective control of cortical and subcortical networks by central thalamus. eLife 4:1–27. doi:10.7554/eLife.09215.001

Lu J, Sherman D, Devor M, Saper CB. 2006. A putative flip–flop switch for control of REM sleep. Nature 441:589–594. doi:10.1038/nature04767

Mankad S, Michailidis G. 2013. Structural and functional discovery in dynamic networks with non-negative matrix factorization. Phys Rev E 88:042812. doi:10.1103/PhysRevE.88.042812

Moore JT, Chen J, Han B, Meng QC, Veasey SC, Beck SG, Kelz MB. 2012. Direct Activation of Sleep-Promoting VLPO Neurons by Volatile Anesthetics Contributes to Anesthetic Hypnosis. Current Biology 22:2008–2016. doi:10.1016/j.cub.2012.08.042

Moruzzi G, Magoun HW. 1949. Brain stem reticular formation and activation of the EEG. Electroencephalography and Clinical Neurophysiology 1:455–473. doi:10.1016/0013-4694(49)90219-9

Nelson LE, Guo TZ, Lu J, Saper CB, Franks NP, Maze M. 2002. The sedative component of anesthesia is mediated by GABAA receptors in an endogenous sleep pathway. Nat Neurosci 5:979–984. doi:10.1038/nn913

Nir Y, Staba RJ, Andrillon T, Vyazovskiy VV, Cirelli C, Fried I, Tononi G. 2011. Regional Slow Waves and Spindles in Human Sleep. Neuron 70:153–169. doi:10.1016/j.neuron.2011.02.043

Ohiorhenuan IE, Mechler F, Purpura KP, Schmid AM, Hu Q, Victor JD. 2010. Sparse coding and high-order correlations in fine-scale cortical networks. Nature 466:617–621. doi:10.1038/nature09178

Owen AB, Perry PO. 2009. Bi-cross-validation of the SVD and the nonnegative matrix factorization. Ann Appl Stat 3. doi:10.1214/08-AOAS227

Pan B, Zucker RS. 2009. A General Model of Synaptic Transmission and Short-Term Plasticity. Neuron 62:539–554. doi:10.1016/j.neuron.2009.03.025

Patel SR, Ballesteros JJ, Ahmed OJ, Huang P, Briscoe J, Eskandar EN, Ishizawa Y. 2020. Dynamics of recovery from anaesthesia-induced unconsciousness across primate neocortex. Brain 143:833–843. doi:10.1093/brain/awaa017

Poulet JFA, Petersen CCH. 2008. Internal brain state regulates membrane potential synchrony in barrel cortex of behaving mice. Nature 454:881–885. doi:10.1038/nature07150

Purpura DP. 1968. Role of synaptic inhibition in synchronization of thalamocortical activity. Prog Brain Res 22:107–122. doi:10.1016/s0079-6123(08)63499-8

Quairiaux C, Megevand P, Kiss JZ, Michel CM. 2011. Functional Development of Large-Scale Sensorimotor Cortical Networks in the Brain. Journal of Neuroscience 31:9574–9584. doi:10.1523/JNEUROSCI.5995-10.2011

Raz A, Grady SM, Krause BM, Uhlrich DJ, Manning KA, Banks MI. 2014. Preferential effect of isoflurane on top-down vs. bottom-up pathways in sensory cortex. Frontiers in Systems Neuroscience 8:191. doi:10.3389/fnsys.2014.00191

Reitz SL, Wasilczuk AZ, Beh GH, Proekt A, Kelz MB. 2021. Activation of Preoptic Tachykinin 1 Neurons Promotes Wakefulness over Sleep and Volatile Anesthetic-Induced Unconsciousness. Current Biology 31:394-405.e4. doi:10.1016/j.cub.2020.10.050

Saczynski JS, Marcantonio ER, Quach L, Fong TG, Gross A, Inouye SK, Jones RN. 2012. Cognitive Trajectories after Postoperative Delirium. New England Journal of Medicine 367:30–39. doi:10.1056/NEJMoa1112923

Sanchez-Vives MV, McCormick DA. 2000. Cellular and network mechanisms of rhytmic recurrent activity in neocortex. Nature Neuroscience 3:1027–1034. doi:10.1038/79848

Saper CB, Fuller PM, Pedersen NP, Lu J, Scammell TE. 2010. Sleep State Switching. Neuron 68:1023–1042. doi:10.1016/j.neuron.2010.11.032

Schiff ND. 2008. Central thalamic contributions to arousal regulation and neurological disorders of consciousness. Annals of the New York Academy of Sciences 1129:105–118. doi:10.1196/annals.1417.029

Schneidman E, Berry MJ, Segev R, Bialek W. 2006. Weak pairwise correlations imply strongly correlated network states in a neural population. Nature 440:1007–1012. doi:10.1038/nature04701

Self MW, Kerkoerle T van, Supèr H, Roelfsema PR. 2013. Distinct roles of the cortical layers of area V1 in figure-ground segregation. Current biology 23:2121–2129. doi:10.1016/j.cub.2013.09.013

Stecker MM, Cheung AT, Pochettino A, Kent GP, Patterson T, Weiss SJ, Bavaria JE. 2001. Deep hypothermic circulatory arrest: I. Effects of cooling on electroencephalogram and evoked potentials. The Annals of Thoracic Surgery 71:14–21. doi:10.1016/S0003-4975(00)01592-7

Steriade M. 2004. Acetylcholine systems and rhythmic activities during the waking--sleep cycle. Prog Brain Res 145:179–196. doi:10.1016/S0079-6123(03)45013-9

Steriade M, Amzica F. 1998. Slow sleep oscillation, rhythmic K-complexes, and their paroxysmal developments. Journal of Sleep Research 7:30–35. doi:10.1046/j.1365-2869.7.s1.4.x

Steriade M, Amzica F, Contreras D. 1994. Cortical and thalamic cellular correlates of electroencephalographic burst-suppression. Electroencephalography and Clinical Neurophysiology 90:1–16. doi:10.1016/0013-4694(94)90108-2

Steriade M, Mccormick DA, Sejnowski TJ. 1993a. Thalamocortical Oscillations in the Sleeping and Aroused Brain. Science 262:679–685.

Steriade M, Nunez A, Amzica F. 1993b. A novel slow (< 1 Hz) oscillation of neocortical neurons in vivo: depolarizing and hyperpolarizing components. J Neurosci 13:3252–3265. doi:10.1523/JNEUROSCI.13-08-03252.1993

Strogatz SH. 2015. BifurcationsNonlinear Dynamics and Chaos: With Applications to Physics, Biology, Chemistry, and Engineering. Boca Raton: CRC Press. pp. 45–94.

Tang A, Jackson D, Hobbs J, Chen W, Smith JL, Patel H, Prieto A, Petrusca D, Grivich MI, Sher A, Hottowy P, Dabrowski W, Litke AM, Beggs JM. 2008. A Maximum Entropy Model Applied to Spatial and Temporal Correlations from Cortical Networks In Vitro. J Neurosci 28:505–518. doi:10.1523/JNEUROSCI.3359-07.2008

Timofeev I, Grenier F, Steriade M. 2004. Contribution of Intrinsic Neuronal Factors in the Generation of Cortically Driven Electrographic Seizures. Journal of Neurophysiology 92:1133–1143. doi:10.1152/jn.00523.2003

Timofeev I, Steriade M. 1996. Low-frequency rhythms in the thalamus of intact-cortex and decorticated cats. J Neurophysiol 76:4152–4168. doi:10.1152/jn.1996.76.6.4152

Tkačik G, Marre O, Amodei D, Schneidman E, Bialek W, Ii MJB. 2014. Searching for Collective Behavior in a Large Network of Sensory Neurons. PLOS Computational Biology 10:e1003408. doi:10.1371/journal.pcbi.1003408

Tkačik G, Schneidman E, Berry II MJ, Bialek W. 2006. Ising models for networks of real neurons. arXiv:q-bio/0611072.

Vazey EM, Aston-Jones G. 2014. Designer receptor manipulations reveal a role of the locus coeruleus noradrenergic system in isoflurane general anesthesia. Proc Natl Acad Sci U S A 111:3859–3864. doi:10.1073/pnas.1310025111

Victor JD, Drover JD, Conte MM, Schiff ND. 2011. Mean-field modeling of thalamocortical dynamics and a model-driven approach to EEG analysis. PNAS 108:15631–15638. doi:10.1073/pnas.1012168108

Vyazovskiy VV, Olcese U, Hanlon EC, Nir Y, Cirelli C, Tononi G. 2011. Local sleep in awake rats. Nature 472:443–447. doi:10.1038/nature10009

Warnaby CE, Sleigh JW, Hight D, Jbabdi S, Tracey I. 2017. Investigation of Slow-wave Activity Saturation during Surgical Anesthesia Reveals a Signature of Neural Inertia in Humans. Anesthesiology 127:645–657. doi:10.1097/ALN.0000000000001759

Witten IH, Frank E, Hall MA. 2011. Data mining: practical machine learning tools and techniques, 3rd ed. ed, Morgan Kaufmann series in data management systems. Burlington, MA: Morgan Kaufmann.

Yu S, Huang D, Singer W, Nikolic D. 2008. A small world of neuronal synchrony. Cereb Cortex 18:2891–2901. doi:10.1093/cercor/bhn047

Zhang Z, Ferretti V, Güntan İ, Moro A, Steinberg EA, Ye Z, Zecharia AY, Yu X, Vyssotski AL, Brickley SG, Yustos R, Pillidge ZE, Harding EC, Wisden W, Franks NP. 2015. Neuronal ensembles sufficient for recovery sleep and the sedative actions of α2 adrenergic agonists. Nat Neurosci 18:553–561. doi:10.1038/nn.3957

Zhou W, Cheung K, Kyu S, Wang L, Guan Z, Kurien PA, Bickler PE, Jan LY. 2018. Activation of orexin system facilitates anesthesia emergence and pain control. Proc Natl Acad Sci USA 115:E10740–E10747. doi:10.1073/pnas.1808622115

Zilles K, Palomero-Gallagher N. 2017. Multiple Transmitter Receptors in Regions and Layers of the Human Cerebral Cortex. Front Neuroanat 11:78. doi:10.3389/fnana.2017.00078

